# Structure of Blm10:13S proteasome intermediate reveals parallel assembly pathways for the proteasome core particle

**DOI:** 10.1101/2024.11.04.621988

**Authors:** Mandeep Kaur, Xiang Chen, Stella Y. Lee, Tyler M. Weaver, Bret D. Freudenthal, Kylie J. Walters, Jeroen Roelofs

## Abstract

Proteasomes are formed by chaperone-assisted assembly of core particles (CPs) and regulatory particles (RPs). The CP chaperone dimer Pba1/Pba2 binds early to proteasome subunits, and is thought to be replaced by Blm10 to form Blm10:CP, which promotes ATP-independent degradation of disordered proteins. Here, we present evidence of distinct parallel assembly pathways for CP by solving five cryo-EM structures including a Blm10:13S pre-assembly intermediate. Our data conflict with the current model of Blm10 and Pba1/Pba2 sequential activity in a single assembly pathway, as we find their CP binding is mutually exclusive and both are present on early and late assembly intermediates. CP affinity for Pba1/Pba2 is reduced during maturation, promoting Pba1/Pba2 release. We find Blm10 undergoes no such affinity switch, suggesting this pathway predominantly yields mature Blm10-bound CP. Altogether, our findings conflict with the current paradigm of sequential CP binding to instead indicate parallel assembly pathways by Pba1/Pba2 and Blm10.

## Introduction

The proteasome is an essential cellular complex responsible for selectively degrading proteins into short peptides. The 26S and 30S proteasome complexes, which degrade ubiquitinated proteins, consist of a 750 kDa core particle (CP) capped by a 700 kDa regulatory particle (RP) at either (26S) or both (30S) ends. The RP contains receptors to recognize ubiquitinated proteins, deubiquitinating enzymes to remove ubiquitin, and an AAA-ATPase ring that facilitates unfolding and translocation of substrates into the CP ^1–3^. The proteolytic activity occurs within the CP, which adopts a hollow cylindrical shape, with two β-rings sandwiched between two α-rings. Each ring is formed from seven homologous subunits (α1-α7 or β1-β7), with β1, β2, and β5 containing caspase-like, trypsin-like, and chymotrypsin-like proteolytic activities, respectively ^4–6^. The CP cylinder has a gated entrance at either end that is closed or opened, as controlled by the N-terminal tails of the α subunits ^7^. While degradative functions for CP by itself have been reported, it generally requires regulators to induce gate opening and other substrate-processing activities. In addition to the RP, other regulatory factors can cap the CP. A common factor amongst many of these regulators is a C-terminal HbYX motif (i.e., an antepenultimate hydrophobic residue, a penultimate tyrosine, and any C-terminal residue) that docks into inter-subunit pockets of the CP α-ring and contributes to gate opening.

Accurate and efficient proteasome assembly is critical for maintaining proteasome levels, the reduction of which causes proteotoxic stress and contributes to the development of neurodegenerative disorders and cancer ^8–13^. The assembly of the CP is assisted by three proteasome-specific assembly chaperone complexes: Pba1/Pba2 (PAC1/PAC2), Pba3/Pba4 (PAC3/PAC4), and Ump1 (POMP). Pba1/Pba2 and Pba3/Pba4 promote α subunit ring formation; the HbYX motifs of Pba1 and Pba2 dock into the CP α5/α6 and α6/α7 pockets, respectively ^14^, while at the opposite side, Pba3/Pba4 promotes α3 incorporation into the ring ^15^. The β-subunits associate sequentially with the α-ring and incorporation of β4, which follows β2 and β3, causes the steric expulsion of Pba3/Pba4 ^16^. Pba1/Pba2 remains bound to the immature complex until maturation, as enabled by their positioning at the outer surface of the α-ring ^14,17^.

Ump1 interacts with both α and β subunits at the same side of the α-ring as Pba3/Pba4 and contributes to the recruitment of the additional β subunits ^14,18,19^. β7 is the last subunit to get incorporated (creating the half CP ⍺_1-7_ β_1-7_), and its propeptide C-terminus contributes to the dimerization of two half CPs into a pre-holocomplex ^20^. The final step in maturation of CP, the cleavage of the propeptides from β1, β2, β5-7, and the degradation of Ump1, reduces the affinity of Pba1/Pba2, freeing the now mature CP to bind RP.

Blm10 (PA200 or PSME4)-bound mature CP functions in the ATP-independent degradation of short peptides and proteins with intrinsically disordered regions, contributing to spermatogenesis, gene regulation by degrading acetylated histones, and mitochondrial health ^21–25^. More recently in humans, this regulator was discovered to be upregulated in tumors, causing attenuated antigenic diversity and impacting immunotherapy ^26^. Blm10 levels are typically substoichiometric compared to CP complexes, with overexpression yielding increased Blm10-bound CP complexes at the expense of almost all 26S and 30S complexes otherwise present ^27^. Blm10 also associates with CP assembly intermediates ^20,28,29^. Deletion of Blm10 in yeast strains with proteasome assembly defects exacerbates growth phenotypes ^20,30^, suggesting that Blm10 contributes to proteasome assembly. Several models show Blm10 interacting with CP after Pba1/Pba2 dissociates and prior to RP binding ^31,32^; however, there is limited experimental support for this model. It is unclear what role Blm10 plays when bound to assembly intermediates and how cells regulate levels of Blm10-bound CP versus RP-bound CP. While structures have been reported for both Blm10:CP and PA200:CP ^33–35^, there is no structure of Blm10 bound to an assembly intermediate, limiting our ability to understand the changes that occur during assembly and the role Blm10-assembly intermediates play in proteasome formation.

Here, we solved cryo-electron microscopy (cryo-EM) structures for five CP assembly intermediates. These structures indicate that Blm10 and Pba1 binding to CP intermediates is mutually exclusive. As we can detect both Blm10 and Pba1/Pba2 on early-stage and late-stage intermediates, our data support a model whereby Blm10 and Pba1/Pba2 function in parallel CP assembly pathways instead of sequentially acting in one common assembly pathway. Finally, the lack of interactions between Blm10 and Ump1 or β propeptides suggest that unlike Pba1/Pba2, Blm10 remains bound throughout CP maturation.

## Results

### Blm10:13S and Blm10:α-ring structures

To evaluate how Blm10 contributes to proteasome assembly, we purified CP assembly intermediates by using a yeast strain with the CP chaperone Ump1 C-terminally TAP-tagged ^17^. We also deleted the Pba1-encoding gene PBA1 to omit the abundant assembly intermediate complexes 13S (onwards referred to as Pba1/2:13S), which includes α1-α7, β2-β4, and Pba1/Pba2, and pre-15S, which contains α1-α7, β2-β6, and Pba1/Pba2 ^14^. The purified proteasomes were prepared for cryo-EM data collection on a Krios G3i microscope equipped with a Gatan3 camera, as described in Methods.

Following 2D classification and refinement, we solved the structures of two Blm10-containing protein complexes that constituted 44% of particles to <3 Å resolution and three additional complexes that lack Blm10 (Fig. 1 and Extended Data Fig. 1).

**Figure 1.**
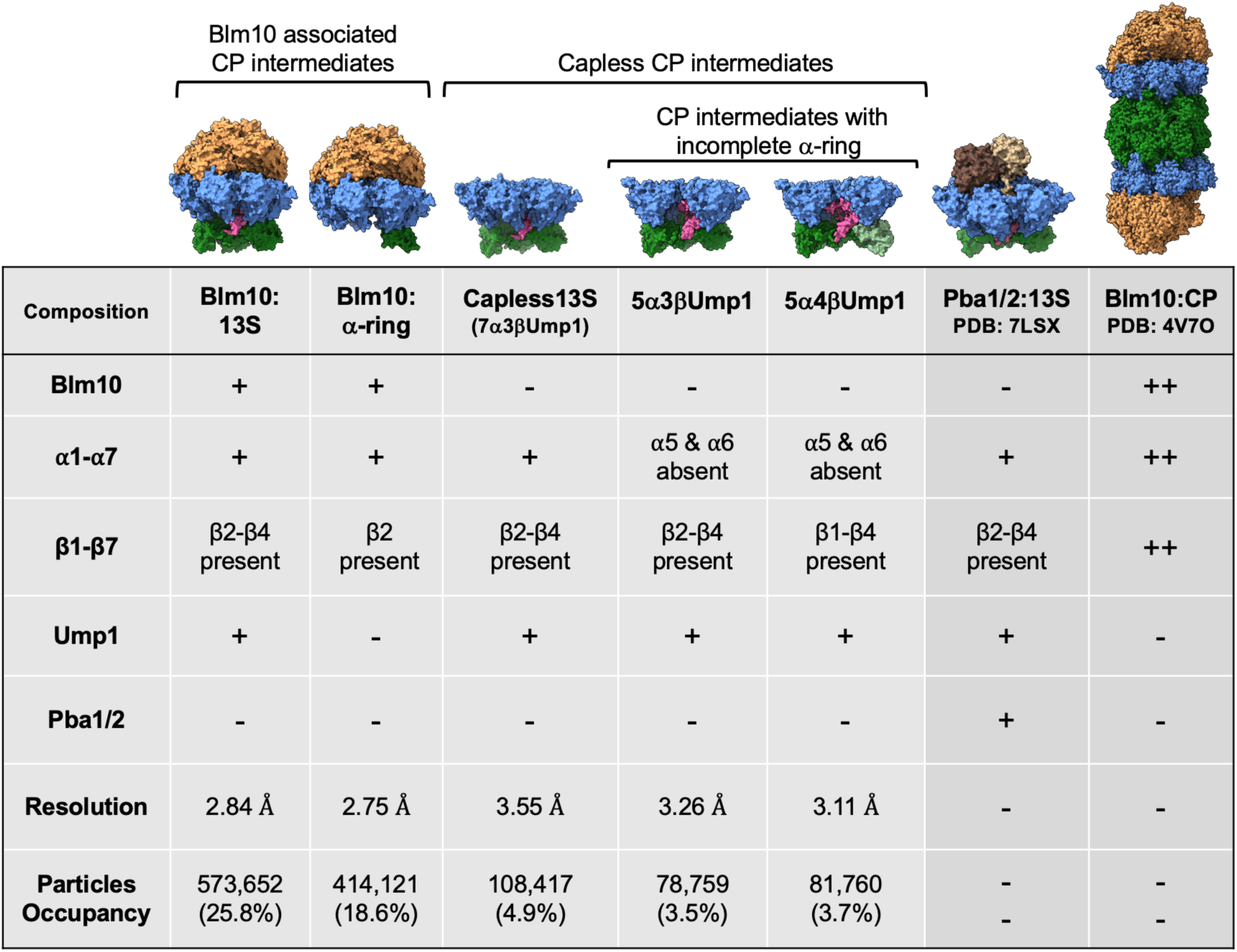
Comparison of the subunit composition of the CP assembly complexes. Cryo-EM structures of five different CP assembly intermediates (Blm10 associated and capless) obtained in this study with their corresponding subunit composition, resolution and particle occupancy are compared to the existing structures of Pba1/2:13S (a.k.a. 13S, PDB: 7LSX) and Blm10:CP (PDB: 4V7O). + and ++ indicates presence of one and two of the respective subunits.

Blm10:13S, which includes α1-α7, β2/β3/β4, and Ump1, was the most abundant complex and resolved to 2.84 Å resolution (Fig. 2a-b and Extended Data Fig. 2a-e). This complex resembles an octopus with three stubby arms formed by the CP β subunits. Blm10 forms a hollow dome with its C-terminal HbYX motif docked into the α5/α6 pocket as the α5 and α6 N-terminal ends insert into Blm10 ( Fig. 2b expanded view and Fig. 2c).

**Figure 2.**
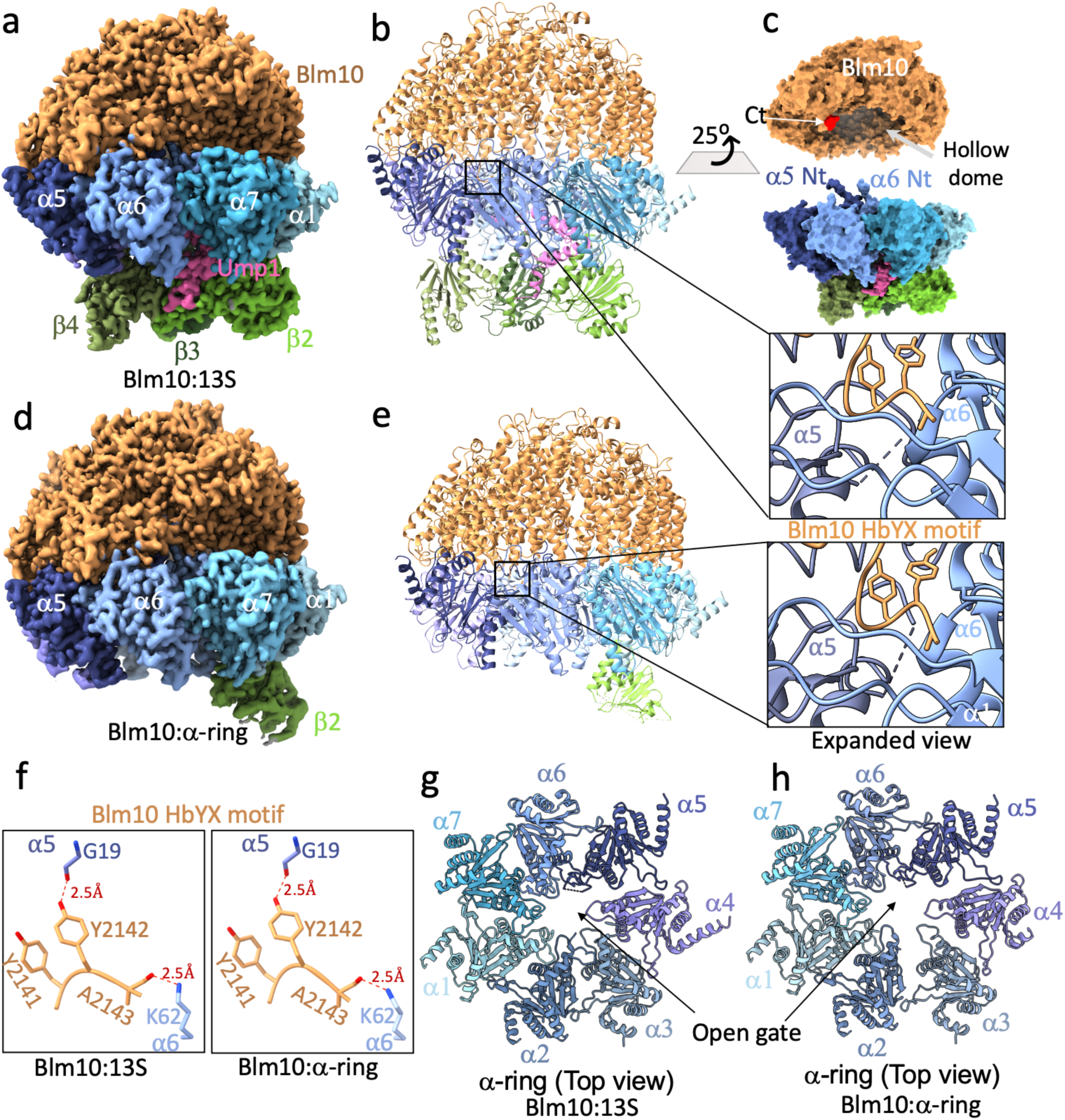
Structure of Blm10:13S and Blm10:α-ring. **a-b,** Cryo-EM density map (2.84 Å, **a**) and corresponding ribbon diagram (**b**) of a Blm10:13S complex that includes the complete α-ring (α1 α7, shades of blue) and β-subunits β2, β3, and β4 (shades of green), Blm10 (orange), and Ump1 (pink). Expanded view shows the position of C-terminus (HbYX motif) of Blm10 (orange) inserted into the α5/α6 inter-subunit pocket. **c,** Surface diagram of Blm10:13S where Blm10 was separated and rotated 25° to reveal termini critical for the interaction**. “**Ct” represents the C-terminus of Blm10 and the HbYX motif is shown in orange. **d-e**, same as **a-b** for Blm10:α-ring (2.75 Å). (**f**) Expanded view of the two important electrostatic interactions (dashed blue line) of the C-terminal HbYX motif of Blm10 (orange) with α5 and α6 (dark and light blue respectively) in Blm10:13S (left panel) and Blm10:α-ring (right panel). (**g,h**) Top view of the α-ring with Blm10 hidden to illustrate the open-gate CP conformation. In (**b**), (**e**), (**g**) and (**h**), a dashed line connects residue endpoints where no density is observed.

A complex containing Blm10, α1-α7, and β2 that lacks Ump1, referred to below as Blm10:α-ring, was resolved to 2.75 Å resolution (Fig. 2d-e and Extended Data Fig. 2f-h). We rationalized that tagged Ump1, by which the complex was isolated, was lost during sample handling. Ump1 becomes increasingly structured by interacting with the β subunits as they are added during CP assembly ^18^ and a complex with fewer β subunits is likely more prone to Ump1 loss. β2 has been previously proposed to be the first β subunit added to the α-ring ^14,18^, a model consistent with β2 being the only β subunit present in our Blm10:α-ring structure (Fig. 2d).

When Blm10 is bound to mature CP, there is one hydrogen bond interaction and one salt-bridge between the HbYX motif and the α5α6 pocket that are of critical importance for association between Blm10 and the α-ring, as their disruption abrogates Blm10 binding to CP ^21,36^. These are hydrogen bond between the sidechain of Blm10 Tyr2142 and α5 Gly19 and a salt bridge between the C-terminal carboxylate of Blm10 (Ala2143) and Lys62 of α6. These interactions aid in opening of the CP gate by repositioning Pro17 of α5 ^29^ and are also observed in Blm10:13S (Fig. 2f, left panel) and Blm10:α-ring (Fig. 2F, right panel). These newly resolved Blm10-bound structures also have an opened CP gate (Fig. 2g-h). Thus, the binding of Blm10 to assembly intermediates utilizes features shared with Blm10 bound to mature CP.

### Capless13S and partial α-ring complexes

Of the three complexes that lack Blm10, one complex, which we refer to as capless13S, contains a complete α-ring, Ump1, and β2/β3/β4 (Fig. 3a and Extended Data Fig. 3a-c). Considering proteasomes can assemble without Blm10 and Pba1/Pba2, this complex likely reflects a genuine 13S-like assembly intermediate, albeit it is possible that RP was lost during sample handling. Surprisingly, in the capless13S complex, the N-termini of the α subunits are disordered, and the ring has an opened gate without the docking of an HbYX motif (Fig. 3a, right panel). Potentially, this complex cycles between an open and closed state, as has been observed for CP ^37^ despite X-ray crystallography only capturing a closed conformation of CP. Nonetheless, no closed form of capless13S was observed and it is possible that a closed state cannot be achieved with only β2/β3/β4 present.

**Figure 3.**
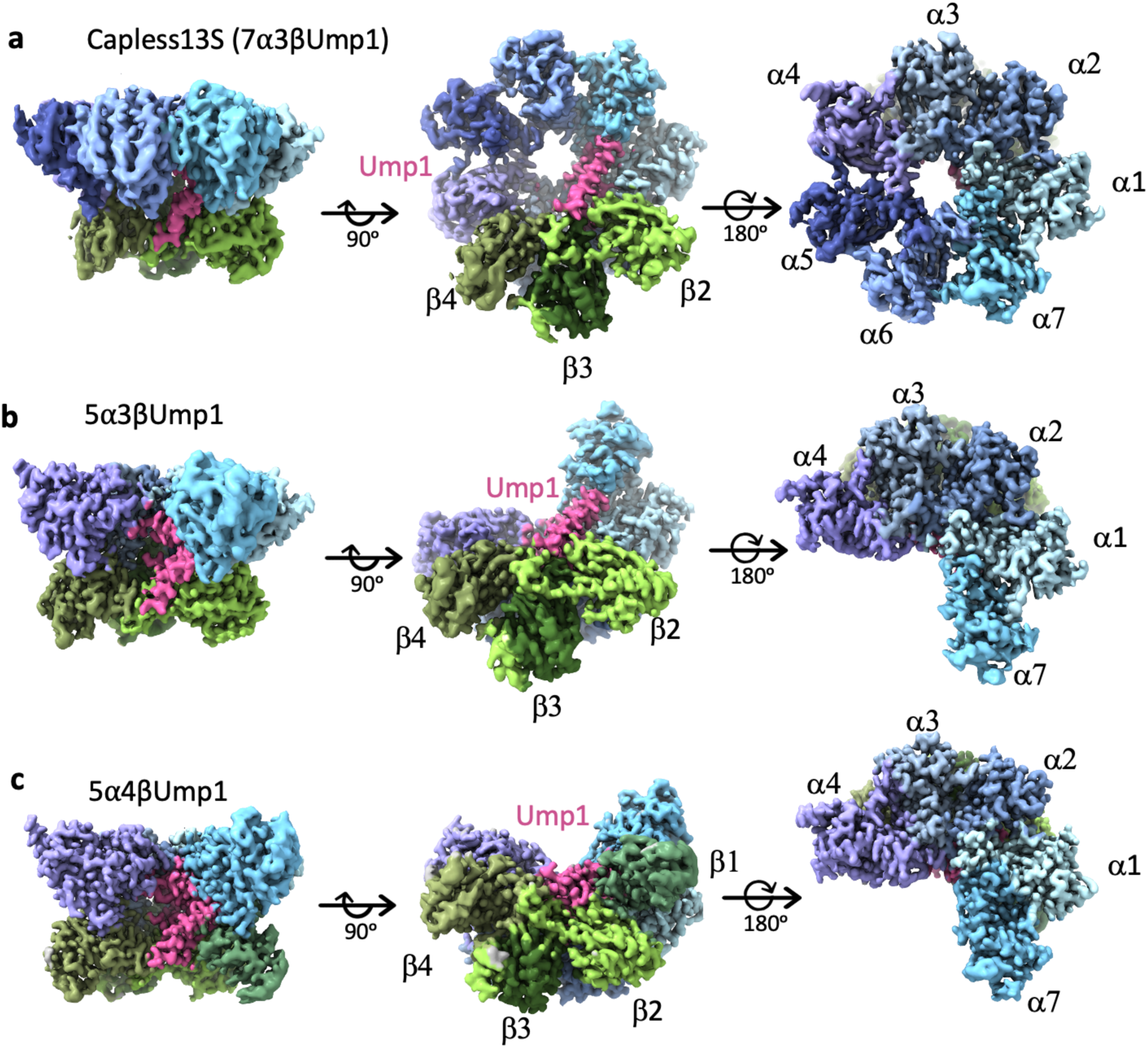
Structure of the capless13S, and partial α-ring complexes. **a-c,** Cryo-EM density maps of capless13S (7α3βUmp1, resolution 3.55 Å), 5α3βUmp1 (resolution 3.26 Å) and 5α4βUmp1 (resolution 3.11 Å) as seen from a side view (left panels), bottom view (middle panels) and top view (right panel). α subunits are represented in shades of blue and purple, β subunits in shades of green, and Ump1 is pink.

The other complexes, 5α3βUmp1 and 5α4βUmp1, consist of α1/α2/α3/α4/α7 and Ump1, with either β2/β3/β4 (Fig. 3b, Extended Data Fig. 3d-f) or β1/β2/β3/β4 (Fig. 3c, Extended Data Fig. 3g-i), respectively. The identification of complexes that have β subunits while lacking a complete α-ring was unexpected as the α-ring is generally thought to form first ^18,38^. However, it is consistent with our previous observations ^17^ and such intermediates have been observed in mutant strains ^39^. Since ring structures are generally stable ^40^ and the missing subunits are those with which Pba1/Pba2 or Blm10 interact (α5 and α6), the presence of these complexes suggests that Pba1/Pba2 and Blm10 facilitate the incorporation of α5 and α6 into the α-ring. There is no structural indication that 5α3βUmp1 and 5α4βUmp1 represent dead-end products because all subunits are correctly ordered in these intermediates. Thus, at least in this *pba1*Δ background, assembly does not always proceed by formation of the α-ring first.

### Blm10:13S and Ump1-mediated conformational changes

During CP assembly, Ump1 directly interacts with multiple α and β subunits and appears to contribute to propeptide cleavage ^19^. Furthermore, Ump1 is degraded as part of CP maturation. Thus, the presence or absence of Ump1 presents as a logical driver of structural changes that occur upon CP maturation. Comparison of Ump1-containing Blm10:13S to Ump1-lacking Blm10:α-ring or Blm10:CP revealed several differences including a conformational change in the Blm10 dome. The Blm10 dome in Blm10:CP contains a large opening of 14 x 23.2 Å that is situated laterally above α2 and α3 (Fig. 4a, left panel). This opening is speculated to allow entry and subsequent degradation of intrinsically disordered proteins ^29^. In Blm10:13S this opening is substantially smaller, at 6.3 x 11 Å (Fig. 4a, middle panel). This constriction is caused by Blm10 loops spanning Asp155-Asp166 and Gly221-Val238, which are ordered in the Blm10:13S structure, but missing in the Blm10:CP structure (Fig. 4b middle and left panel and Extended Data Fig. 4a) ^29^. There are multiple hydrogen bonds and hydrophobic interactions in Blm10:13S that stabilize the Blm10 loops spanning Asp155-Asp166 and Gly221-Val238 (Extended Data Fig. 4b). Since the smaller opening in Blm10:13S would cause steric hindrance to the entry for an unfolded polypeptide, we refer to this configuration as in a closed state. The changes in the dome are accompanied by changes at the interface between α3 and loops of Blm10 that connect helix 49 (H49) to helix 50 (H50) and helix 46 (H46) to helix 47 (H47). Here, hydrophobic and three hydrogen bond interactions are observed in Blm10:13S that are absent in Blm10:CP (Extended Data Fig. 4c and d). Looking at the Blm10:⍺-ring structure (lacking Ump1), the dome opening was intermediate in size, at 12.7 x 17.5 Å (Fig. 4a and b, right panel). The Blm10 loops Asp155-Asp166 and Lys229-Val238 are observable, but Gly221-Gly228, which was observed in the Blm10:13S structure, is not (Fig. 4b right panel). In all, these data suggests that the Blm10 dome opening can change from a closed state to an open state, enabling potential allosteric regulation of Blm10-dependent protein degradation.

**Figure 4.**
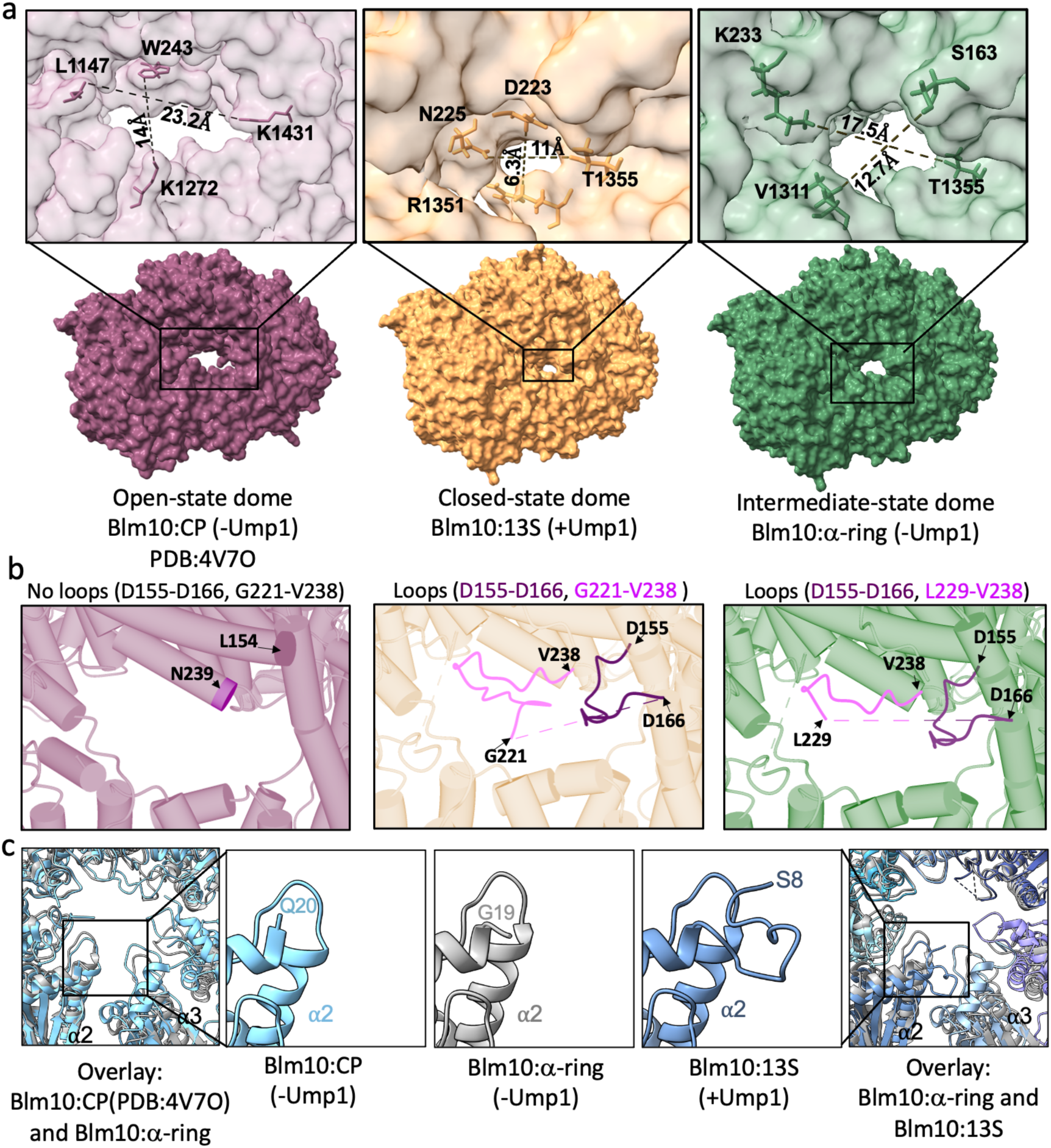
Ump1 mediated changes in the Blm10:CP interaction. **a**, Comparison of the diameter of the largest Blm10 dome opening in Blm10:CP (magenta), Blm10:13S (orange) and Blm10:α-ring (green). Corresponding expanded views are shown on top with dimensions. **b,** Expanded view of Blm10 dome opening with α helices represented by tubes showing no loops between residue Leu154 and Asn239 in Blm10:CP (left panel), how the dome opening is covered by loops Asp155-Asp166 (dark pink) and Gly221-Val238 (light pink) in Blm10:13S (middle panel), and loops Asp155-Asp166 (dark pink) and Lys229-Val238 (light pink) for Blm10:α-ring (right panel)**. c,** Superposition of the α-ring of Blm10:CP (cyan) and Blm10:α-ring (grey) (left-most), and panels showing the expanded view of the first resolved residues of the N-terminus of α2 in Blm10:CP, Blm10:α-ring, and Blm10:13S, as indicated. The far right panel shows the superposition of the α-ring of Blm10:13S (in shades of blue) and Blm10:α-ring (grey).

Another difference uniquely correlated with the presence of Ump1 is the status of the α2 N-terminal residues. These residues are disordered in Blm10:CP, based on the absence of electron density for the first 19 residues (Fig. 4c). However, in Blm10:13S, residues from Ser8 onwards are resolved in the structure and partially fill the CP gate (Fig. 4c and Extended Data Fig. 5). The α2 N-terminus is disordered in Blm10:α-ring, more closely resembling Blm10:CP (where Ump1 is absent) than Blm10:13S (Fig. 4c). This comparison suggests that the maturation of CP, which is accompanied by degradation of Ump1 causes increased flexibility of the α2 N-terminal residues (accompanied by a more opened CP gate) and transition from a closed Blm10 dome to an opened state, which likely activates the complex. CP assembly and the degradation of Ump1 is likely to induce additional conformational changes. For example, superimposing the β-subunits of Blm10:13S and Blm10:CP, there are steric clashes that need to be resolved. This requires a rearrangement of β2 and β3 to prevent steric clashes with Ump1 and the β2 pro-peptide (Extended Data Fig. 6).

### Comparison of Blm10:13S to Pba1/2:13S rationalizes differences in specificity for immature CP

The HbYX motifs of Blm10 and Pba1 occupy the same α5/α6 CP pocket (Fig. 2b, expanded view) ^14,41^. Nevertheless, aligning Blm10:13S with Pba1/2:13S highlights differences in the CP gate opening (Fig. 5a and b) and the α2-α5 side of the α-ring in Blm10:13S is shifted down towards the β-ring, with the largest displacement for α4 (Fig. 5c). In Blm10:13S there is a loss in electron density for α2 Met1 to Phe7, α3 Met1 to Phe13, and α4 Met1 to Phe18 resulting in an open gate (Fig 5a, left and middle panel). An open gate, however, might not enable activity for this complex because the Blm10 dome is closed (Fig. 4a, middle panel). In Pba1/2:13S, despite the HbYX motif docking, most of the N-terminal residues of α2, α3 and α4 are ordered due to interactions with the Pba1 N-terminal tail, which plugs the gate (Fig. 5a and b, middle and right panels)^14^.

**Figure 5.**
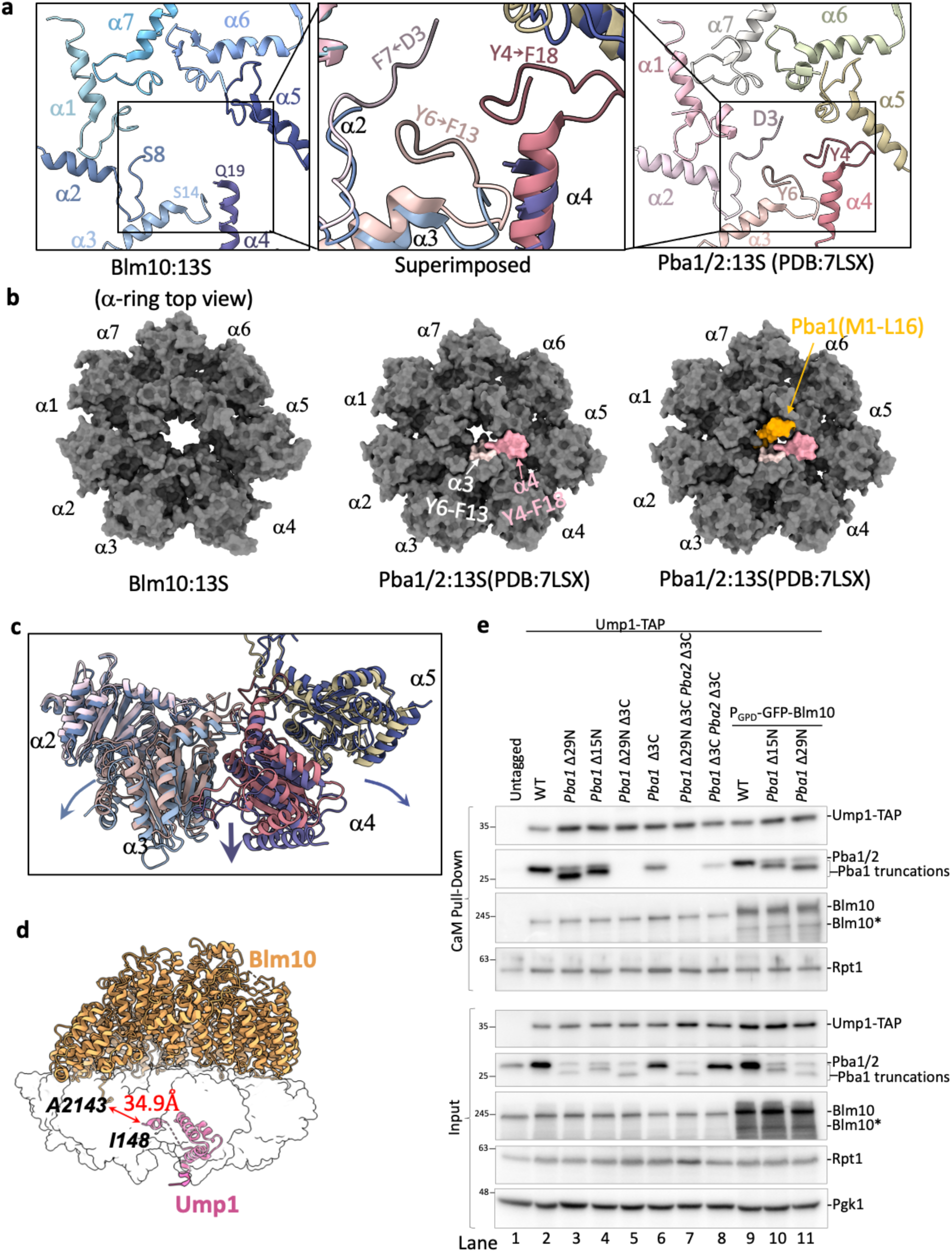
Differences in Blm10:13S vs Pba1/2:13S structure. **a**, N-terminal tails of α1-α7 subunits indicating the first resolved residues in Blm10:13S (left panel), Pba1/2:13S (right panel) and an expanded view of superimposed Blm10:13S and Pba1/2:13S (middle panel). The latter shows the additional resolved residues of α2 ( D3-F7), α3 (Y6-F13) and α4 (Y4-F18) that contribute to the restricted gate in Pba1/2:13S. **b,** Top view of the α-ring showing the extent of gate opening upon binding of Blm10 (left) or Pba1/Pba2 (middle). Light and dark pink surfaces in the middle and right panels represent the α3 and α4 residues of Pba1/2:13S, respectively, that are missing from Blm10:13S. The right panel is identical to the middle panel, but includes the Pba1 N-terminal tail (orange). **c,** The superimposed α-ring of Blm10:13S (shades of blue and purple) and Pba1/2:13S (shades of pink and green) showing only α2 - α5 subunits. Arrows indicate the downward displacement of the subunits, with largest shift for α4. **d,** Blm10:13S structure with Blm10 and Ump1 displayed as ribbons and the other subunits as transparent white surfaces. The distance between the C-terminal residue of Blm10 (A2143) and Ump1 (I148) is displayed and indicated with a red solid line. **e,** Pba1 N-terminus contributes to its affinity for immature CP. Strains with N- or C-terminal truncations of Pba1 were lysed and Ump1 containing complexes were purified using Calmodulin resin. Following lysis, samples were separated by SDS-PAGE, transferred to a PVDF membrane and immunoblotted for indicated proteins. Top panel shows immunoblot analysis of Ump1-TAP Calmodulin (CaM) pull-down from the specified yeast strains. Bottom panel shows the total cell lysate (input) from the respective strains. Blm10* is a Blm10 specific band likely reflecting a breakdown product.

Pba1’s N-terminal tail also interacts with Ump1 and the β5 propeptide ^14^. For Blm10, the closest residue to Ump1 is Ala2143, which is 34.9 Å away (Fig. 5d), indicating there are no direct interactions between Blm10 and Ump1. Despite this difference, Ump1 is positioned almost identically in both 13S structures (Extended Data Fig. 7a), indicating that Pba1 and Blm10 do not influence its positioning. The lack of interactions between Blm10 and intermediate complex specific components (i.e., the propeptides or Ump1) suggests that Blm10 is not subject to dramatic changes in affinity during CP maturation as Ump1 or the propeptides are degraded. Consistent with this interpretation, Blm10 remains bound when Ump1 is absent, as revealed in the Blm10:α-ring structure (Fig. 1) and Blm10-bound CP complexes. In all, our data provide a structural rationale for the presence of Blm10 on assembly intermediates as well as mature CP, while Pba1/Pba2 solely associates with intermediates ^17,27,41^.

To determine whether the Pba1 N-terminal tail plays a role in regulating the amount of Blm10 versus Pba1/Pba2 that is bound to 13S, we aimed to disrupt the Ump1-Pba1 interaction by mutating Ump1 residues Arg93 and Glu89, which interact with the Pba1 N-terminus (Extended Data Fig. 7b). However, we did not see any difference in Pba1 binding to Ump1, perhaps because the propeptides also interact with Pba1. Therefore, we next tested the impact of N-terminal truncations of Pba1. Using small deviations compared to previous efforts ^42^, we were able to readily detect Pba2 and N-terminally truncated Pba1 variants (either 15 residues, *pba1*Δ15N, or 29 residues, *pba1*Δ29N) in the cell lysates, albeit at lower levels compared to full length Pba1 (Fig. 5e, bottom, lane 3 and 4 compared to lane 2). While the N-terminal truncations by themselves did not abrogate Pba1/Pba2 binding to assembly intermediates, we detected an impact on binding when combined with truncations of the HbYX motif of Pba1 alone (Pba1Δ3C) or in combination with Pba2 (Pba1Δ3C Pba2Δ3C) (Fig. 5e). Strikingly, the deletion of both Pba1 and Pba2 HbYX motifs showed more residual binding to 13S than Pba1 with its N- and C-terminus truncated (*pba1* β29N β3C, Fig. 5e, lane 8 versus lane 5). In all, these data show the importance of the Pba1 N-terminus for binding to intermediates.

Nevertheless, altering Pba1/Pba2 interaction with 13S did not alter the amount of copurified Blm10 (Fig. 5e) and we wondered if the substoichiometric levels of Blm10 relative to CP in yeast could obscure any putative competition. Indeed, upon overexpression of Blm10, the *pba1*Δ15N and *pba1*Δ29N showed a larger reduction in Pba1/Pba2 on Ump1-containing assembly intermediates as compared to full length Pba1 (Fig. 5e, lane 2, 3, and 4 versus 9, 10, and 11). Thus, the N-terminal tail of Pba1 contributes to the high affinity of Pba1/Pba2 for assembly intermediates and helps to prevent any excess Blm10 from displacing Pba1/Pba2 from assembly intermediates.

In all, our analyses highlight a crucial difference between Blm10 and Pba1/Pba2 regarding their interaction within the 13S complexes. For Pba1/Pba2, there are interactions with Ump1 and β-subunit propeptides that contribute to an affinity switch whereby Pba1/Pba2 loses affinity upon maturation of the CP (i.e. when Ump1 and propeptides get degraded). Blm10 lacks these interactions and does not display an affinity switch to release it upon CP maturation. Instead, Blm10 remains bound to CP upon maturation to form Blm10:CP complexes.

### Parallel pathways of CP assembly

The binding of Blm10 and Pba1/Pba2 to an α-ring is mutually exclusive and they have previously been proposed to bind sequentially ^31,32^. Sequential binding can occur by Blm10 binding after Pba1/Pba2 expulsion (Fig. 6a, top panel, path *i*), or alternatively, with Blm10 binding first (Fig. 6a, top panel, path *ii*). The latter model is inconsistent with retention of Blm10 on mature CP. Yet, Pba1 is found on late assembly complexes, such as Ump1-containing 13S, 15S, and pre-holoenzyme complexes, which is inconsistent with the former model. Therefore, we postulated that instead of a sequential pathway, a parallel assembly pathway is more consistent with the experimental data and protein structures (Fig. 6a, lower panel). If Blm10 and Pba1/Pba2 act in parallel (i.e., different) pathways of assembly, Blm10 and Pba1/Pba2 would both be present at both early and late assembly intermediates. In a sequential pathway, at least one of them should be absent early or late in assembly.

**Figure 6.**
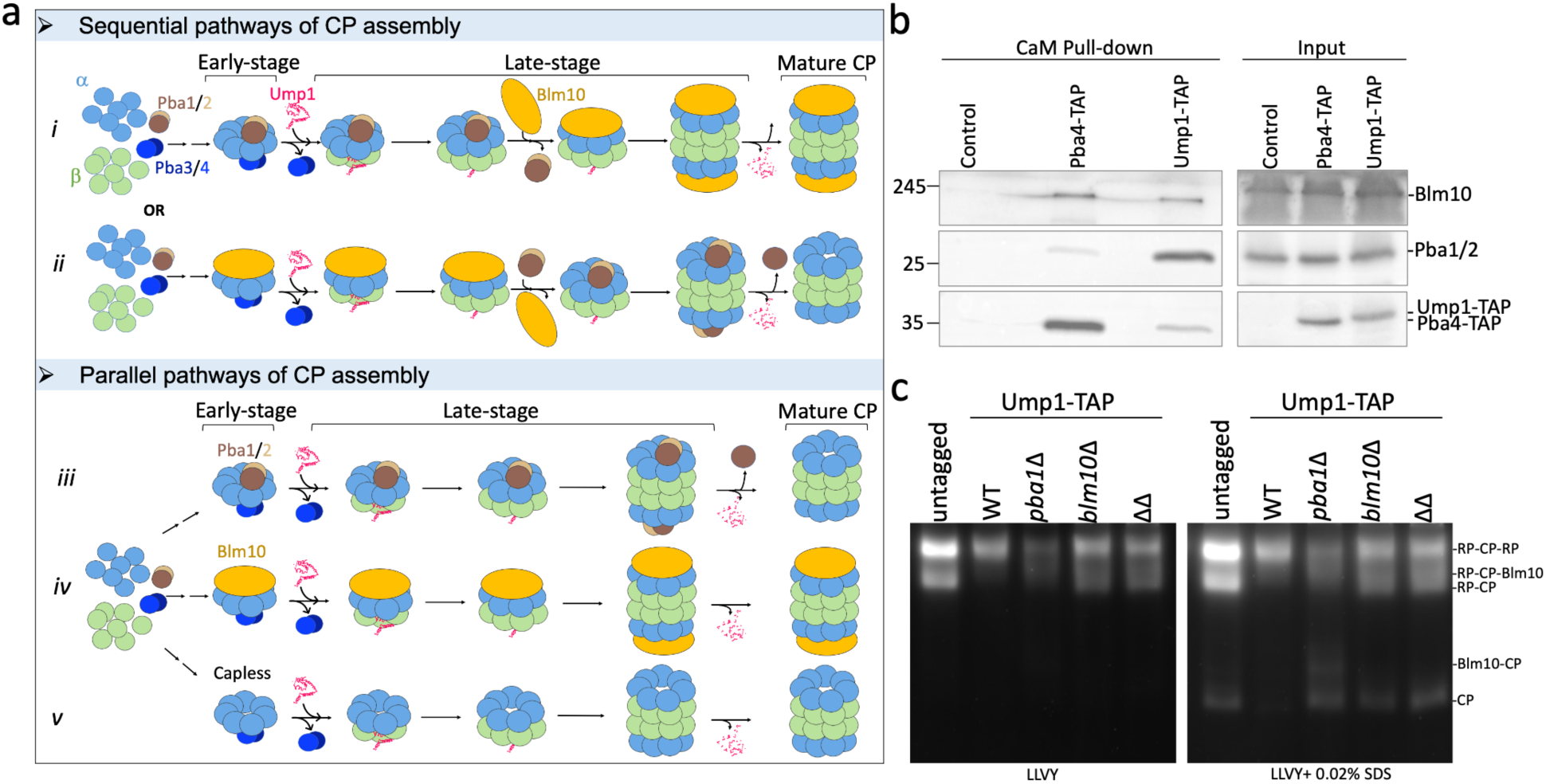
Parallel pathways of proteasome assembly. **a**, Conceptual model of CP assembly via sequential or parallel assembly pathways. The 20S assembly starts with the formation of α-ring assisted by Pba1/Pba2 and Pba3/Pba4 dimers. The β subunits are sequentially added with the help of Ump1 which replaces the Pba3/Pba4 dimer. Together with pro-peptides, Ump1 helps in assembling the half CP rings into a pre-holocomplex. In a sequential pathway (top panel), either Pba1/Pba2 binds in the early stages of assembly and gets replaced by Blm10 in the late stages (*path i*) or vice versa (*path ii*), ultimately resulting in mature free CP (details in text). In a parallel pathway (bottom panel), CP follows three parallel pathways, (*path iii*) Pba1/Pba2 associated, (*path iv*) Blm10 associated and (*path v*) with no cap (RP associated, potentially). **b**, Calmodulin (CaM) based pull down assay to detect the presence of Pba1/Pba2 and Blm10 in the early-stage (Pba4 pull-down) and late stage (Ump1 pull-down) in the left panel. Blm10 and Pba1/Pba2 are present in early as well as late-stage CP assembly intermediates. The right panel shows the total cell lysates (input) from the respective yeast strains. The wild-type strain with no TAP tag served as a negative control. **c**, Native gel showing proteasome activity in total cell lysates of different strains. The left panel shows activity without SDS and right panel shows activity after incubating the gel in 0.02% SDS.

Early steps in the CP assembly pathway can be distinguished from later steps by the different chaperones that associate with the complex; Pba3/Pba4 binds the ⍺-ring during early assembly steps (Fig. 6a, dark blue), whereas Ump1 binds to the complex at later steps (Fig. 6a red). A steric clash between Pba4 and Ump1 ensures that upon incorporation of Ump1, Pba3/Pba4 is expelled from the assembly intermediate, providing a clear definition of early versus late assembly intermediates; while a recent cryo-EM study revealed one structure with PAC3/PAC4 and POMP within the same complex amongst a series of human CP assembly intermediates ^18^, it should be noted that these structures were purified from heterologously expressed human proteins. All studies looking at endogenous complexes have not identified this complex, suggesting that it is a short-lived, low abundance complex under physiological conditions. In both yeast and humans, Pba3/Pba4 leaves early during CP assembly, Ump1 arrives later and remains bound until a mature CP has been produced ^18,38^.

Performing pulldowns from Pba4-tagged (early) and Ump1-tagged (late) yeast strains, we detected Pba1/Pba2 in both purifications (Fig. 6b). Blm10, which we know binds to mature CP as well as the 13S complex that contains Ump1 (our Blm10:13S structure), also copurified with early- and late-stage intermediates (Fig. 6b, Pba4-TAP and Ump1-Tap lanes, respectively). Thus, both Blm10 and Pba1 are present in early as well as late stages of CP assembly, which is consistent with parallel assembly pathways. As we also solved the structure of capless13S (i.e., 13S-like complex without Blm10 or Pba1/Pba2 (Fig. 1, 3a)), there are likely three parallel assembly pathways to form ½ CP: Blm10-dependent, Pba1/Pba2-dependent, and a pathway that is independent of both (potentially assisted by RP ^20^). The latter pathway must exist, as proteasomes can still form in strains deleted of both PBA1 and BLM10 (Fig. 6c).

Whether the formation of mature CP follows preferred combinations of ½ CPs and how this assembly relates to the formation of hybrid proteasome complexes (such as Blm10:CP:RP) remains to be determined.

## Discussion

The accurate and efficient assembly of proteasomes depends on proteasome-specific chaperones, including the CP chaperones Pba1, Pba2, Pba3, Pba4, and Ump1. Here, we report the cryo-EM structure of the Blm10:13S assembly intermediate to find mutually exclusive binding of Blm10 and Pba1/Pba2 to the 13S based on overlapping interaction surfaces at the α-ring and common binding of their HbYX motifs to the α5/α6 pocket. This raises questions of how and when Blm10 binds to the α-ring and whether Pba1/Pba2 remains bound until the CP is fully assembled. Our data show that Blm10 and Pba1/Pba2 are present both early in assembly (Pba4-based pulldown) and late in assembly (Ump1-based pulldown), which is inconsistent with the prevailing model that Blm10 binds after Pba1/Pba2 in the assembly pathway ^31,32^. Instead, we propose a model whereby Blm10:13S, Pba1/2:13S, and a capless13S are each intermediates in parallel pathways of CP assembly (Fig. 6a, lower panel).

### Functionality of parallel assembly pathways

In the model of parallel assembly pathways, there are numerous advantages that would provide the cell benefits and adaptability. First, parallel pathways can provide robustness to the CP assembly process, by having another pathway available when one is compromised. Indeed, upon deletion of PBA1, the upregulation of proteasome subunits (via rpn4 stabilization) becomes critical and Blm10 or RP might substitute for Pba1/Pba2 function ^20^. Furthermore, the *blm10*β *pba1*β cells are still capable of producing CP and 26S proteasomes. Thus, when needed, there is redundancy in CP assembly.

Another function of parallel pathways could be that it allows for the formation of different proteasome complexes, whose ratio can be controlled at the step of assembly. One striking difference between the Pba1/Pba2-dependent assembly pathway and Blm10-dependent pathway is the different fates of the chaperone following CP maturation. For Pba1/Pba2, there is a dramatic drop in affinity for the CP when propeptides are removed and Ump1 is degraded. This affinity switch ensures Pba1/Pba2 is tightly bound during assembly, but not associated with mature CP. Thus, Pba1/Pba2’s sole purpose is to allow for efficient CP assembly and once formed, CP is available for association with RP, PA28 family members (absent from yeast), or Blm10 (PA200). In the Blm10:13S structure, Blm10 is 34 Å away from Ump1 and does not have any direct interactions with the features of CP that are indicative of the immature form (Ump1 or propeptides). As such, the degradation of Ump1 upon CP maturation does not impact the affinity of Blm10 for the α-ring. Consistent with this model, we solved a structure of Blm10:α-ring, which has Blm10 bound in the absence of Ump1. Thus, it appears that the function of Pba1/Pba2 dependent assembly is the formation of CP, while the Blm10 dependent assembly pathway leads to the formation of Blm10-bound CP complexes. Since Blm10 and the human ortholog PA200 are responsible for the degradation of a variety of substrates ^23,24,36^, this parallel pathway might ensure the presence of a certain amount of Blm10-bound CP complexes in the cell. Interestingly, overexpression of Blm10 in yeast, or the observed high expression of PA200 in mouse testis is accompanied by high cellular levels of Blm10 (PA200)-bound CP and reduced 26S and 30S in cells. It remains to be determined to what extent these are formed from competition between RP and Blm10 for mature CP binding versus being driven by this alternative assembly pathway directly. If the off rates of RP and Blm10 *in vivo* are low, the assembly pathway might be particularly important.

A third putative function of parallel pathways is to deal with misassembled CP intermediates. The existence of proteasome specific assembly chaperones illustrates how complex and crucial the assembly of proteasomes is and reflect the potential of errors in assembly. Since the deletion of BLM10 does not have a significant impact on CP assembly in general, it is intriguing that higher amounts of Blm10-bound CP are reported in mutants that result in CP assembly defects such as *ump1*Δ ^30^ and *PRE4*-CTEΔ ^20^. All these mutants are likely to have reduced fidelity in assembly. Blm10 has also been reported to associate with CPs with an open gate ^30^ and the α3 knockout strain, which has α4-α4 proteasomes with a wider CP gate, also show more Blm10 bound mature CP complexes ^43^. In all, this might reflect a functional link between a non-standard α-ring and Blm10. In humans, this property might have further evolved, where PA200 might be working with a specialized α subunit, such as in spermatozoa where both PA200 and α4s are highly expressed ^23,44^.

### Early assembly

The very early steps of CP assembly, where individual α subunits combine to form an α-ring remain poorly characterized. The structures we solved that lack a complete α-ring, suggest that there is not a strict requirement for the formation of an α-ring before β-subunits are added, as α1, α2, α3, α4, and α7 form a stable intermediate with β2, β3, β4, plus or minus β1. The formation of this complex would not involve the Pba1/Pba2 chaperone, as Pba1/Pba2 intimately binds to α5 and α6 and seems to help with the incorporation of these subunits into the α-ring. We purified α5β3Ump1 and α5β4Ump1 from a *pba1*Δ strain and in wild type cells, these two intermediates are less likely to accumulate. Pba1/Pba2 together with α5 and α6 will quickly be incorporated to form Pba1/2:13S, as this complex has β subunits with propeptides and Ump1, both of which bind the Pba1 N-terminal tail (Extended data Fig. 4). This leaves the possibility open that Blm10 can compete with Pba1/Pba2 for the assembly intermediate particularly when an α-ring is being formed or prior to Ump1 and β subunit incorporation. Consistent with this possibility, a recent cryo-EM structure of a PAC1/PAC2-bound α-ring showed the N-terminal tail, at least of PAC1, was not involved in binding to this intermediate ^45^.

In all, our data show the existence of three parallel pathways of CP assembly and provides more insight in the early assembly process by revealing it is not strictly α-ring first. Our data reveal an additional level of complexity to an already complex process. As we have seen little to no free Blm10 in lysates ^27^, it appears that the Blm10-dependent assembly pathway ensures and enables cells to produce a certain level of Blm10 (or PA200)-bound proteasomes. This allows Blm10 protein expression to control the amount of RP-bound CP verses Blm10-bound CP complexes. This mechanism of regulation has direct relevance in understanding tumors where PA200 is upregulated and influence immunotherapy responses ^26^. It will also aid in understanding the physiological role and regulation of PA200 in spermatogenesis, mitochondrial health and the degradation of acetylated histones ^21–25^.

## Methods

### Yeast strains

All strains were constructed in background *MAT A his3Δ0 leuΔ0 met15Δ0 ura3Δ0* unless mentioned otherwise (Supplementary table 1). Standard PCR based procedures (primers and plasmid templates presented in Supplementary table 2) were applied for the gene deletions. Crispr Cas9 was used to generate Pba1, Pba2, and Ump1 mutations.

### CaM-based pull-down Assay

Indicated yeast strains were grown for 4 hours starting at an OD_600_ = 0.38. Cell lysis was done by cryogrinding. Calmodulin resin (G-Biosciences) was prewashed three times with EDTA-free lysis buffer (50 mM Tris-HCl, pH=7.5, 5 mM MgCl_2_, 1 mM ATP, 2 mM CaCl_2_). The washed resin was incubated with the lysate for 1 hour at 4 °C followed by three washes with wash buffer (50 mM Tris-HCl, pH=7.5, 5 mM MgCl_2_, 1 mM ATP, 2 mM CaCl_2_, 50 mM NaCl). The proteins bound to CaM resin were subjected to SDS-PAGE.

### Western blot analysis

Yeast strains were grown at 30 °C and harvested when the OD_600_ reached 1. Cell pellets equivalent of OD_600_ = 2 were collected by centrifugation, then promptly lysed or stored at −80 °C. Lysis was performed using an alkaline lysis method as previously described or by grinding in Liquid N_2_ ^46,47^. Briefly, the pellets were resuspended in 100 μL distilled water with the addition of 100 μL of 200 mM NaOH, followed by a 5-minute incubation at room temperature. Cell suspensions were pelleted, resuspended in 50 μL SDS-PAGE sample buffer (0.06 M Tris–HCl, pH 6.8, 5% glycerol, 2% SDS, 50 mM DTT, 0.0025% bromophenol blue), boiled at 98 °C for 5 min, and supernatant was collected. Proteins were separated on SDS-PAGE and Western blots were done using antibodies against Blm10 (Rabbit polyclonal; Enzo Life Sciences BML-PW0570), Pba1/2 ^17^, CBP (Mouse; GenScript CBP Tag Antibody cat. No. A01798), Rpt1 (Abcam Aβ22678), Pgk1 (Invitrogen, catalog no.459250). Secondary antibodies conjugated to horseradish peroxidase were from Rockland Immunochemicals. HRP activity was visualized using the Immobilon Forte Western HRP substrate (Millipore). Images were acquired using a Gbox imaging system (Syngene) and captured with GeneSys software.

### Native gel analyses

Yeast cells were harvested at indicated times. Cell pellets equivalent to OD_600_ 50 were centrifuged and lysed by cryo-grinding under non-denaturing conditions as described previously ^46,48^. Protein concentrations were measured using nanodrop. Equal amounts of protein (300 μg) for each strain were loaded on native gel (3.5%). The activity of native proteasome complexes was visualized by an in-gel LLVY-AMC activity assay described previously ^46^. Images were acquired using Gbox imaging system (Syngene).

### Assembly intermediate purification

Proteasome assembly intermediate purification was carried out by two step TAP purification method ^49^. Specific yeast strain with Ump1-TAP tag was grown to a final OD_600_ ∼ 10.0 in a 9 L YPD media. 105 g cell pellet was collected and resuspended in 1.5X pellet volume of lysis buffer (50 mM Tris-HCl, pH 8.0, 1 mM EDTA, Protease inhibitor Roche cOmplete mini tablet) and lyzed by French press (1200 Psi.). IgG beads (MP Biomedical; 1 ml bed volume per 50 g cell pellet were added to the lysate and incubated for 1 hour at 4 °C. IgG Beads were washed with 50 bed volumes of wash buffer 1 (50 mM Tris-HCl, pH 7.5, 100 mM NaCl, 1 mM EDTA) followed by 15 bed-volumes of cleavage buffer (50 mM Tris-HCl, pH 7.5, 5 mM MgCl_2_, 1 mM DDT).

Overnight (at 4 °C) incubation of washed resin in the cleavage buffer with 10 μl TEV protease (Invitrogen) was done to elute the proteasome intermediates with CBP part of the TAP tag still intact with the CP assembly intermediates. To the eluted protein sample (total 5 mg), 2 mM CaCl_2_ was added prior to performing the second affinity purification using CaM (calmodulin SepharoseTM 4B) resin (2 ml slurry). Following 1.5-hour incubation, the resin was washed twice with wash buffer 2 (50 mM Tris-HCl, pH 7.5, 5 mM MgCl_2_, 2 mM CaCl_2_, 50 mM NaCl) and once with wash buffer 3 (50 mM Tris-HCl, pH 7.5, 5 mM MgCl_2_, 50 mM NaCl with no CaCl_2_). Final elution was done using buffer with 2 mM EGTA (50 mM Tris-HCl, pH 7.5, 5 mM MgCl_2_, 2 mM EGTA, 1 mM DDT) after 30 min incubation. The eluate was concentrated to 1.8 mg/ml.

### Cryo-EM sample preparation and data acquisition

3 μL of the purified sample (1.8 or 0.9 mg/mL) were applied on a Quantifoil® R 1.2/1.3 Cu 400 mesh grids (cat. Q4100CR1.3), that had been glow discharged three times (0.42 mBar, 15 mA for 45s.) by using a PELCO easiGlow glow discharge system. Grids were vitrified in 100% liquid ethane using a Leica EM GP 2 plunge freezer (Leica Microsystems) with a blot time of 1 s or 3 s, a blot distance of 43 mm and 80% humidity. Grids were subsequently imaged on a 300 kV Titan Krios microscope (Thermo Fisher Scientific) equipped with a K3 direct electron detector (Gatan) and an energy filter (Gatan). The images were taken at a magnification of 81,000 and the pixel size is 1.068 Å/pixel. 58 frames per movies were acquired for a total dose of 60 elections/Å^2^. 5,235 movies were collected using the EPU program (Thermo Fisher Scientific), with defocus values ranging from −2.5 to −0.9 μm, at 0.2 μm intervals. Details of the data collection and set parameters are listed in Table 1.

**Table 1.** Statistics for cryo-EM analyses.

|  | **Blm10:13S** | **Blm10:α-ring** | **capless13S**<br>(Ump1:α-ring:β2/β3/β4) | **5α3βUmp1**<br>(Ump1:α1/α2/α3/<br>α4/α7:β2/β3/β4) | **5α4βUmp1**<br>(Ump1:α1/α2/α3/<br>α4/α7:β1/β2/β3/β4) |
| --- | --- | --- | --- | --- | --- |
| **Data collection and image processing** |  |  |  |  |  |
| Microscope | Titan Krios |  |  |  |  |
| Camera | K3 |  |  |  |  |
| Magnification | 81,000 |  |  |  |  |
| Voltage (kV) | 300 |  |  |  |  |
| Electron exposure (e<sup>-</sup>/Å<sup>2</sup>) | 60 |  |  |  |  |
| Defocus range (μm) | 0.9 - 2.5 |  |  |  |  |
| Pixel size (Å) | 1.068 |  |  |  |  |
| Symmetry imposed | C1 |  |  |  |  |
| Total micrographs | 5,235 |  |  |  |  |
| Initial particle images | 2,226,993 |  |  |  |  |
| Final particle images | 586,158 | 414,121 | 108,417 | 78,759 | 81,760 |
| Map resolution (Å) | 2.84 | 2.75 | 3.55 | 3.26 | 3.11 |
| FSC threshold | 0.143 | 0.143 | 0.143 | 0.143 | 0.143 |
| Map sharpening *B* factor (Å<sup>2</sup>) | -83.8 | -77.7 | -87.8 | -65.20 | -75.20 |
| EMDB code | 46461 | 46559 | 46570 | 46525 | 46535 |
| **Model building and refinement** |  |  |  |  |  |
| Initial model used (PDB ID) | 7LSX, 4V7O | 7LSX, 4V7O | 7LSX | 7LSX | 7LSX |
| Model resolution (Å) | 2.8 | 2.8 | 3.7 | 3.3 | 2.9 |
| FSC threshold | 0.143 | 0.143 | 0.143 | 0.143 | 0.143 |
| *Model composition* |  |  |  |  |  |
| Non-hydrogen atoms | 67181 | 57789 | 35582 | 28488 | 30875 |
| Amino acid residues | 4230 | 3631 | 2283 | 1822 | 1974 |
| Protein molecules | 12 | 9 | 11 | 9 | 10 |
| *Real-space correlation* |  |  |  |  |  |
| CC (volume) | 0.73 | 0.69 | 0.56 | 0.67 | 0.68 |
| CC (mask) | 0.72 | 0.69 | 0.56 | 0.66 | 0.67 |
| *RMS deviations* |  |  |  |  |  |
| Bond lengths (Å) | 0.014 | 0.014 | 0.014 | 0.014 | 0.014 |
| Bond angles (°) | 1.892 | 1.873 | 1.891 | 1.892 | 1.822 |
| *Validation* |  |  |  |  |  |
| MolProbity score | 0.88 | 0.85 | 0.83 | 0.95 | 0.93 |
| Clash score | 0.01 | 0.00 | 0.00 | 0.00 | 0.00 |
| Rotamers outliers (%) | 0.32 | 0.78 | 0.88 | 1.49 | 1.49 |
| CaBLAM outliers (%) | 3.01 | 3.09 | 1.85 | 2.43 | 2.32 |
| Cβ outliers (%) | 0.00 | 0.00 | 0.00 | 0.00 | 0.00 |
| *Ramachandran plot* |  |  |  |  |  |
| Favored (%) | 94.46 | 94.88 | 95.19 | 95.43 | 95.71 |
| Allowed (%) | 5.54 | 5.12 | 4.81 | 4.57 | 4.29 |
| Outliers (%) | 0.00 | 0.00 | 0.00 | 0.00 | 0.00 |
| PDB code | 9D0T | 9D4C | 9D4U | 9D35 | 9D3I |

### Cryo-EM image processing

All cryo-EM data processing was performed by using cryoSPARC 4.4.1^50^. A flowchart of the data processing is displayed in Extended data Fig. 1 and a summary of cryoEM reconstruction statistics is listed in Table 1. 5235 dose-fractionated movies were gain-reference corrected, aligned, dose-weighted and summed to single-frame micrographs by patch motion correction, followed by constant transfer function (CTF) estimation done by patch CTF estimation. The micrographs were visually inspected and those with broken or thick ice, or CTF resolution > 5 Å were excluded from the stack, leaving 4,786 micrographs. An initial set of particles (589 particles) were manually picked from 11 micrographs and templates were created by selecting 18 classes by running 2D classification. Template picking identified 234,354 particles from 232 micrographs and new templates were created by selecting 28 classes followed by 2D classification. A total of 4,058,518 particles were picked from 4,786 micrographs by template picking and extracted by using a box size of 340 pixels.

Particle stacks were subjected to 2D classification to remove clear false positive particles, carbon edges and junk particles. 2,226,993 particles with various orientations in 2D classes were selected and re-centered. These particles were subjected to one round of ab initio model generation and 3D heterogeneous refinement into 3 classes.

One class of 1,462,486 particles showed presumed Blm10 density (class 1 in Extended data Fig. 1) and was further subjected to two rounds of ab initio model generation and 3D heterogeneous refinement. Particles were sorted and combined into Blm10:13S-like or Blm10:α-ring-like groups based on the presence of presumed β2, β3, β4 and Ump1 density. Each group of particles were subjected to further 2D classification, CTF refinement and non-uniform refinement, resulting in Blm10:13S and Blm10:α-ring maps with overall resolution of 2.84 Å (Blm10:13S with 573,652 particles) and 3.02 Å (Blm10:α-ring with 227,391 particles), respectively, judged by the gold standard Fourier shell correlation (GSFSC) in cryoSPARC (Extended data Fig. 1a). From the first round of 3D heterogeneous refinement, two other classes of particles were detected. One class of 556,986 particles showed a CP-like density map but no presumed Blm10 density (class 2 of Extended data Fig 1). These particles were subjected to similar ab initio model generation, 3D heterogeneous refinement, CTF refinement and non-uniform refinement. Three maps were reconstructed for 7α3βUmp1, 5α3βUmp1, and 5α4βUmp1, with overall resolution of 3.55, 3.26, and 3.11 Å, respectively (Extended data Fig. 1b). The remaining 207,611 particles (class 3 of Extended data Fig. 1) were refined similarly, but the overall resolution of resulting maps was >5.7 Å, so no further analysis was performed on these maps.

In parallel to the first round to 3D heterogeneous refinement, a total of 2,226,993 particles were subjected to a round of ab initio model generation and 3D homogeneous refinement, and resulted in a map of 2.85 Å resolution. One round of unsupervised 3D classification was performed, and particles were sorted into 6 classes. One class with 414,121 particles was different from the other classes and showed map density for only one β subunit. This class of particles was subjected to further CTF refinement and non-uniform refinement, resulting in a reconstructed Blm10:α-ring map with an overall resolution of 2.75 Å (Extended data Fig. 1d). This Blm10:α-ring map was then used for further model building and refinement. All GSFSC and viewing direction distribution plots as well as local resolution maps were generated in cryoSPARC.

### Model building and structure analyses

For atomic model building and refinement, the Blm10 structure from PDB 4V7O and α1– α7, β2–β4, Ump1 structures from PDB 7LSX were fitted as rigid bodies into the Blm10:13S density map by using UCSF ChimeraX ^51^.To assist model interpretation, the refined Blm10:13S map, as well as other refined maps were processed by EMready ^52^. The Blm10:13S structural model was refined by iterative cycles of manual building in Coot (Emsley et al., 2010) and real-space refinement in Phenix ^53^.The geometry and real-space correlation validation were performed by using the phenix.validation_cryoem module in Phenix ^53^. The refined Blm10:13S atomic model was used as an initial model for atomic model building and refinement of Blm10:α-ring, 7α3βUmp1, 5α3βUmp1, and 5α4βUmp1 complexes. For 5α4β:Ump1, the β1 structure of PDB 4V7O was also fit as a rigid body into the density map. A summary of model building and validation statistics is listed in Table 1. Figures were prepared using UCSF ChimeraX.

## Data availability

The cryo-EM maps generated by this study have been deposited into the EMDB database with accession codes EMD-46461 (Blm10:13S), EMD-46559 (Blm10:α-ring), EMD-46570 (7α3βUmp1), EMD-46525 (5α3βUmp1), and EMD-46535 (5α4βUmp1).

The atomic models reported in this paper have been deposited into the PDB with accession codes 9D0T (Blm10:13S), 9D4C (Blm10:α-ring), 9D4U (7α3βUmp1), 9D35 (5α3βUmp1), and 9D3I (5α4βUmp1). The raw micrographs and particle data have been deposited in EMPIAR with accession number EMPIAR-12275. Other data are available from the corresponding authors upon reasonable request.

## Acknowledgements

We like to thank Drs. Lynn Schrag, Sebastian Karuppan, and Lejla Zubcevic for helpful discussions and assistance with negative staining and the Vitrobot use. We thank Gabriel Wooden for help with cloning and Dr. Alicia Burris and all members of the Roelofs lab for feedback on the project and manuscript. We also like to acknowledge the University of Chicago Cryo-EM core for image acquisition and the KUMC cryo-EM facility and directors for their technical assistance and the financial support in the form of a Pilot project award. We acknowledge use of the Frederick Research Computing Environment cluster, NCI at Frederick. This work was supported by grants from the NIH-NIGMS R35-GM149314 (JR), and R35-GM128562 (BDF), and funding from the Intramural Research Program through the Center for Cancer Research, National Cancer Institute, National Institutes of Health (1 ZIA BC011490 to KJW).

---

**Extended Data Fig. 1.**
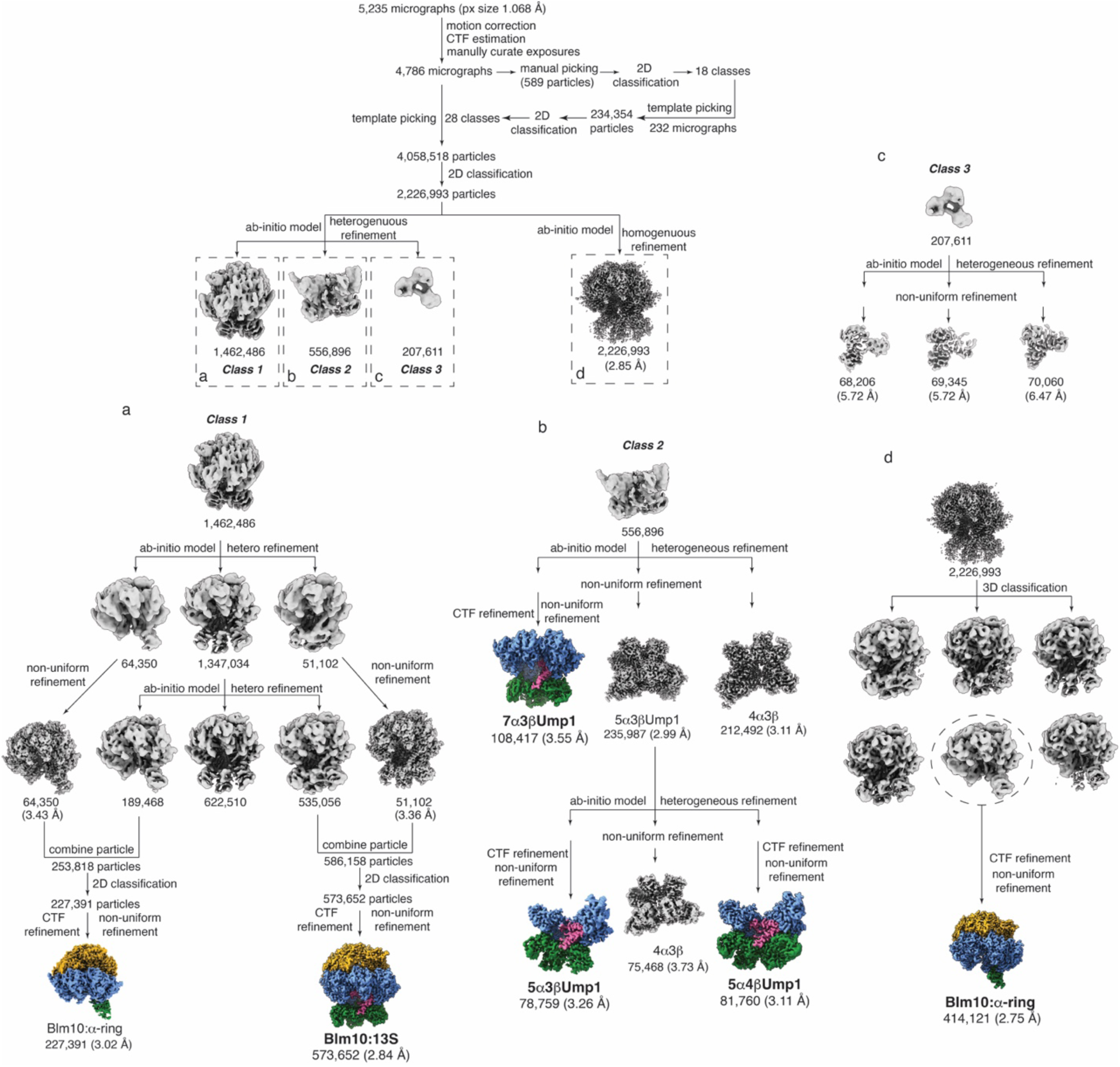
Cryo-EM workflow for complexes purified using an Ump1-affinity tag. Processing scheme for the classification shows reconstructions following both homogeneous and heterogeneous refinement of Blm10:13S, Blm10:α-ring, 7α3βUmp1, 5α3βUmp1, and 5α4βUmp1 complexes. The reconstructed maps are colored as in Fig. 1.

**Extended Data Fig. 2.**
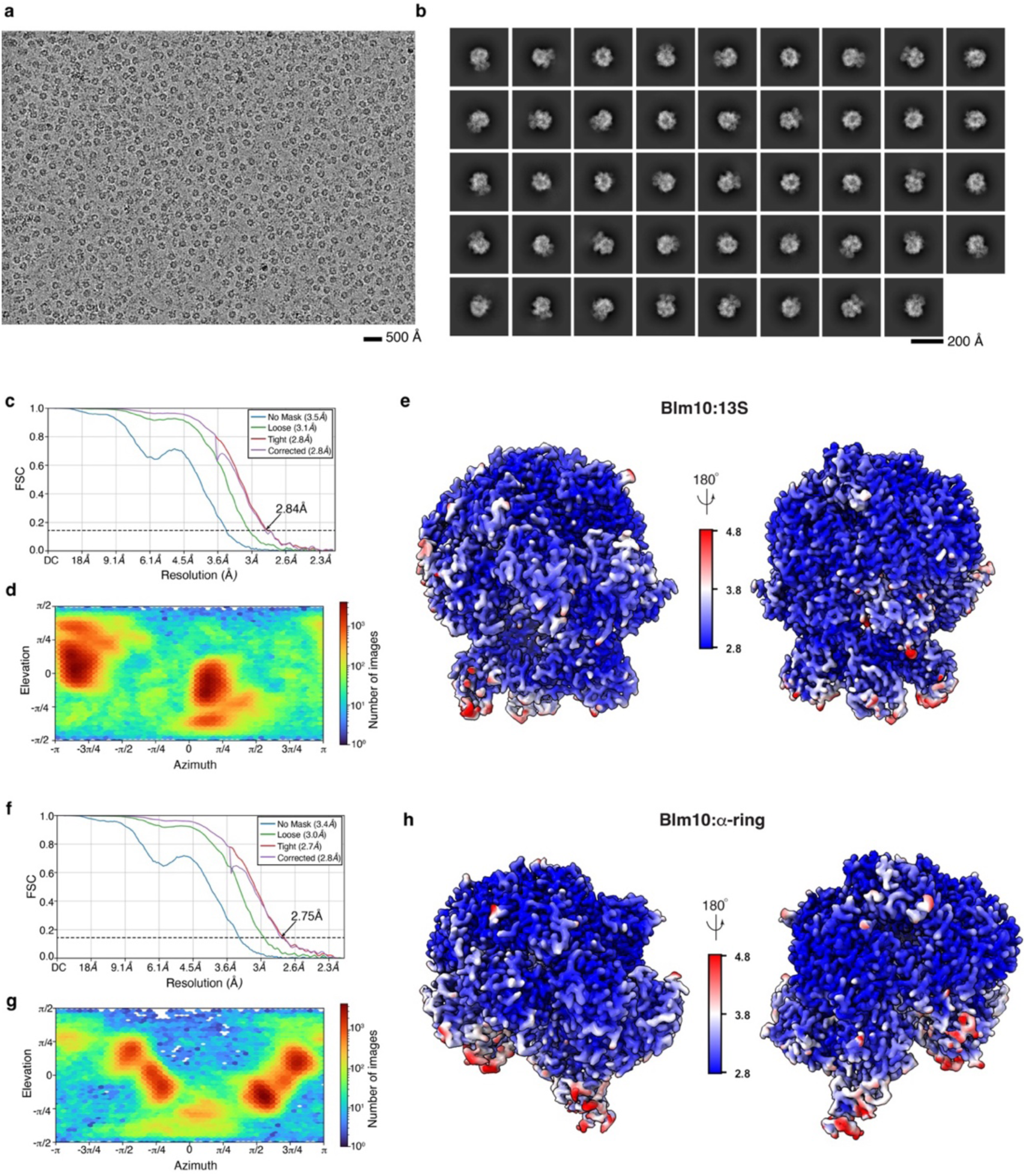
Cryo-EM data analyses for the Blm10:13S and Blm10:α-ring complexes. **a,** Representative cryo-EM micrograph acquired on the Ump1-tagged sample. The data were acquired with a Titan Krios microscope equipped with a Gatan K3 direct detector and energy filter. Scale bar, 500 Å. **b,** Representative 2D class averages computed by cryoSPARC ^50^. Scale bar, 200 Å. **c-h,** Gold standard Fourier Shell Correlation (GSFSC) plots (c and f), viewing direction distribution plots (d and g), and cryo-EM density maps displaying local resolution as calculated by cryoSPARC (e and h) for the refined maps of the Blm10:13S (c, d and e) or Blm10:α-ring (f, g and h) complexes. In c and f, the FSC value of 0.143 is indicated by a black dashed line.

**Extended Data Fig. 3.**
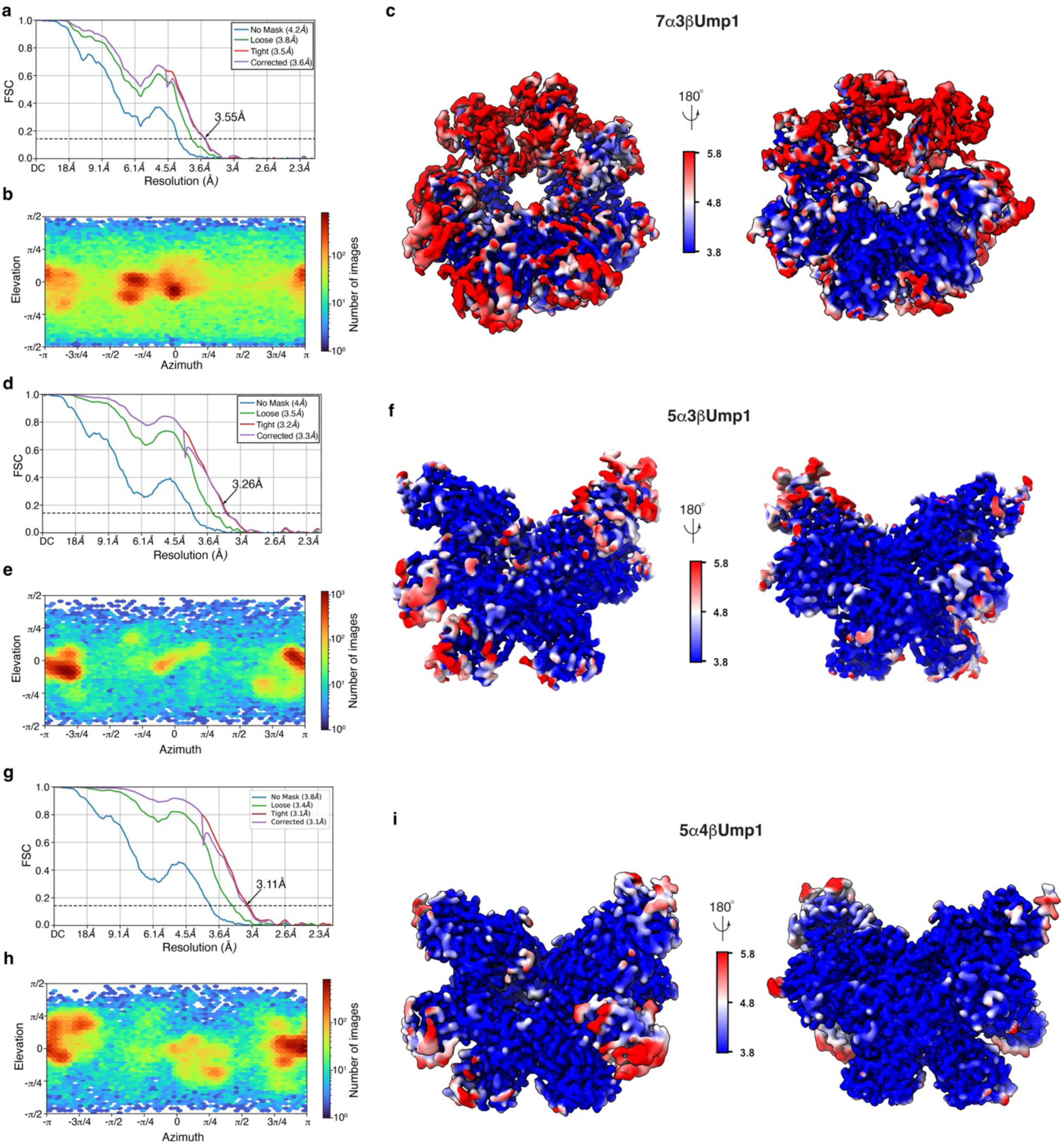
Cryo-EM data analyses for the capless13S and partial α-ring complexes. **a-i,** GSFSC plots (a, d, and g), viewing direction distribution plots (b, e, and h), and cryo-EM density maps displaying local resolution as calculated by cryoSPARC (c, f, and i) for the refined maps of capless13S (a, b, and c), 5α3βUmp1 (d, e, and f), and 5α4βUm (g, h, and i). FSC value of 0.143 is indicated by a black dashed line in a, d, and g.

**Extended Data Fig. 4.**
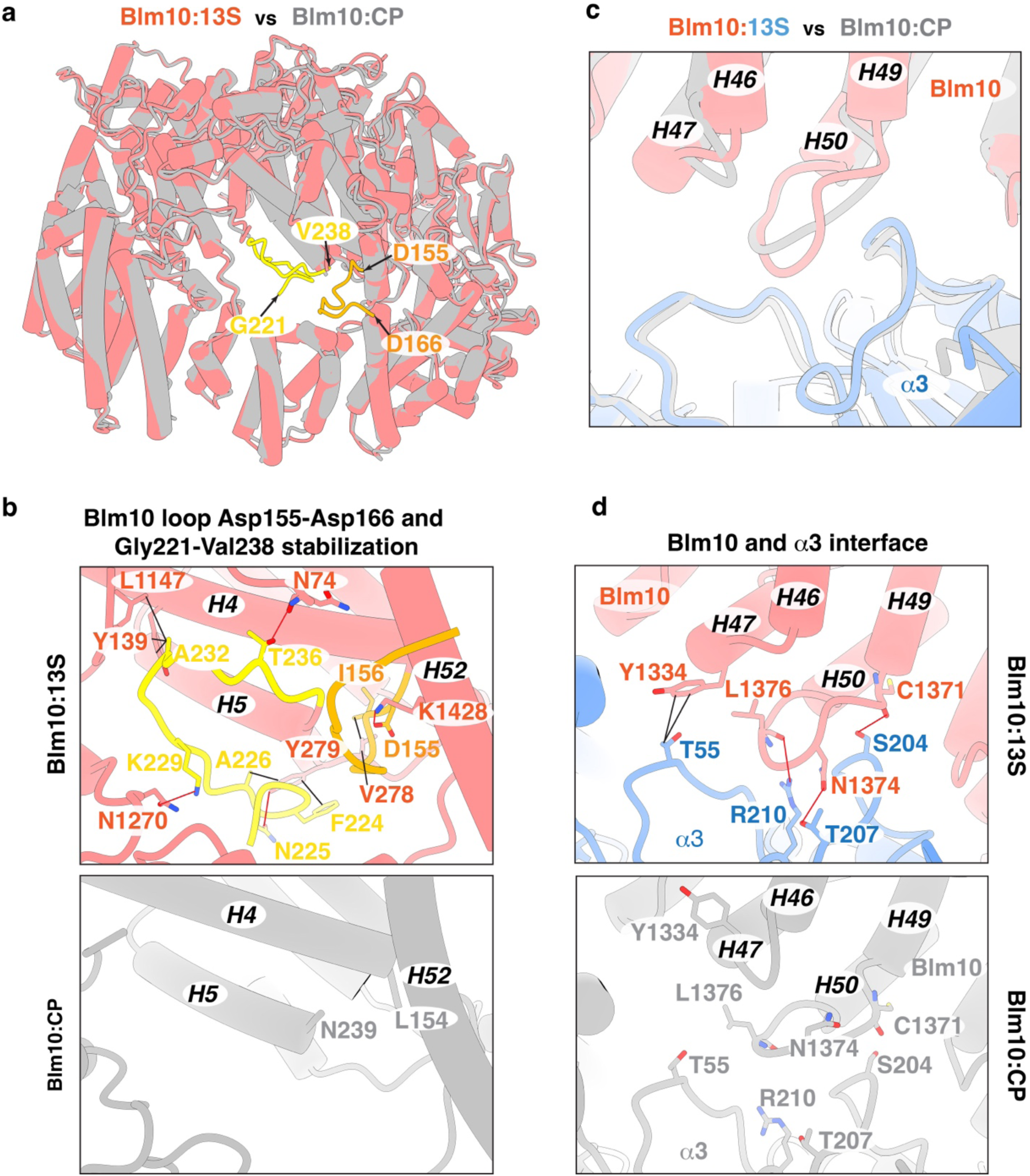
Conformational changes impacting Blm10 dome opening. **a**, Top view of superimposed Blm10 cartoon structures from Blm10:13S (pink) and Blm10:CP (grey). The Blm10 loops identified in Blm10:13S (Asp155-Asp166 and Gly221-Val238) are colored orange and yellow. **b**, Expanded view of Blm10:13S (top) cartoon structure highlighting intramolecular interactions involving the Blm10 loops Asp155-Asp166 (orange) and Gly221-Val238 (yellow). These loops were not observed in the Blm10:CP structure (PDB:4V7O, bottom panel). Blm10 is colored as in (a). The residues involving hydrogen bond or hydrophobic interactions are displayed as sticks with nitrogen and oxygen atoms colored indigo and red. **c**, Expanded view of the superimposed Blm10 (pink):13S (blue) and Blm10: CP (grey, PDB:4V7O) cartoon structures highlighting the rearrangement of the loops connecting Blm10 helices H46 to H47 and H49 to H50. These loops are at the Blm10 (pink):α3 (blue) interface. **d**, Expanded view of the Blm10:13S cartoon structure (top panel) highlighting the additional interactions between Blm10 and α3, which are absent in the Blm10:CP structure (bottom panel). Blm10 and α3 are colored as in (c). Interacting residues in Blm10 and α3 are displayed as sticks with nitrogen, oxygen and sulphur atoms colored indigo, red and yellow respectively. In b and d, hydrogen bonds and hydrophobic interactions are indicated with red and black lines, respectively. In (b)-(d), Blm10 helices H4-H6, H46-H47, H49-H50 and H52 are labeled.

**Extended Data Fig. 5.**
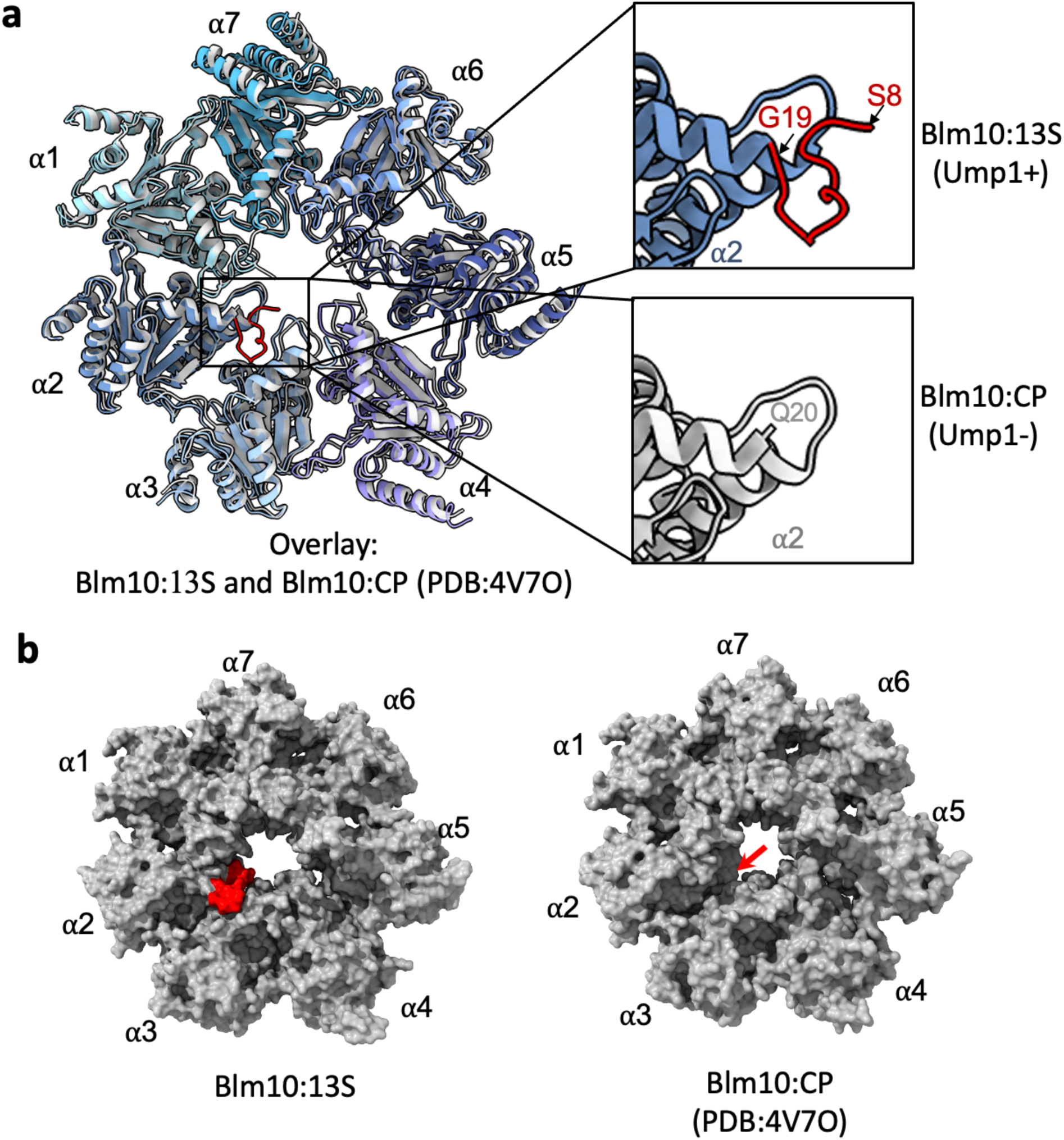
A stabilized α2 N-terminus restricts gate opening in Blm10:13S compared to Blm10:CP. **a,** Top view of the α-ring of Blm10:13S (shades of blue and purple) and Blm10:CP (shades of grey, PDB: 4V7O) with expansions (right) highlighting the more ordered α2 N-terminus (S8-G19, red) of Blm10:13S (top) and the first resolved residue (Q20) of Blm10:CP (grey). **b,** Top view of the molecular surface of the α-ring showing the CP gate architecture for Blm10:13S highlighting the stabilized α2 N-terminus (S8-G19, red, left) and Blm10:CP (right). A red arrow indicates where the N-terminus of α2 is located in Blm10:13S.

**Extended Data Fig. 6.**
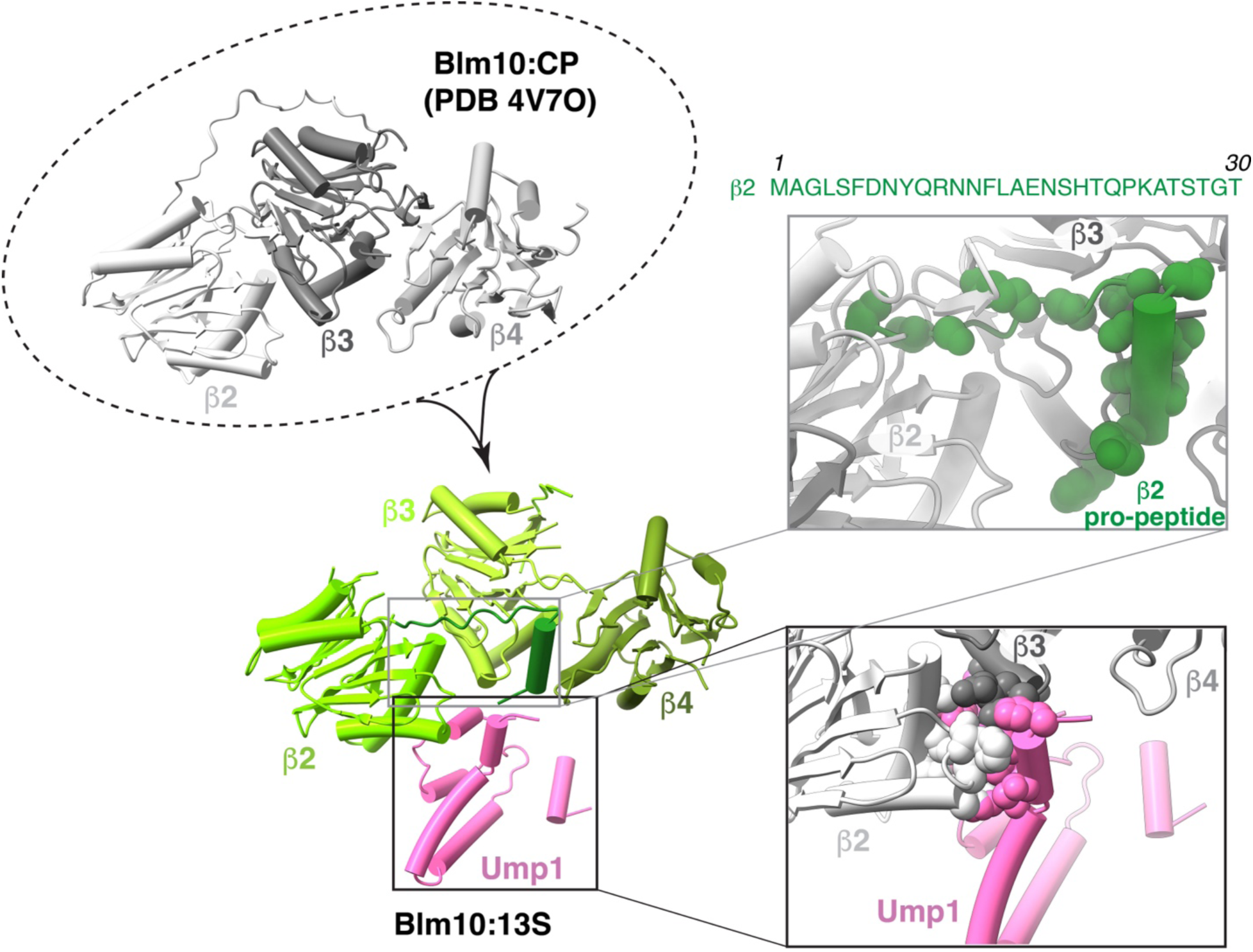
Steric clashes for Ump1 upon aligning Blm10:13S and Blm10:CP indicate conformational changes upon CP maturation. Superposition of the β-ring subunits (β2, β3, and β4) from the Blm10:CP structure (top left panel, shades of grey, PDB: 4V7O) with those in the Blm10:13S structure (bottom left panel, shades of green) and with expansions (top and bottom right panels) showing the rearrangements of β2 and β3 to prevent steric clashes between β2 and Ump1 (pink) or β3 and the β2 pro-peptide (dark green). The residue atoms of β2 pro-peptide (top, right panel), β2 and Ump1 (bottom, right panel) involving steric clashes are displayed as spheres. The amino acid sequence of the β2 pro-peptide is included.

**Extended Data Fig. 7.**
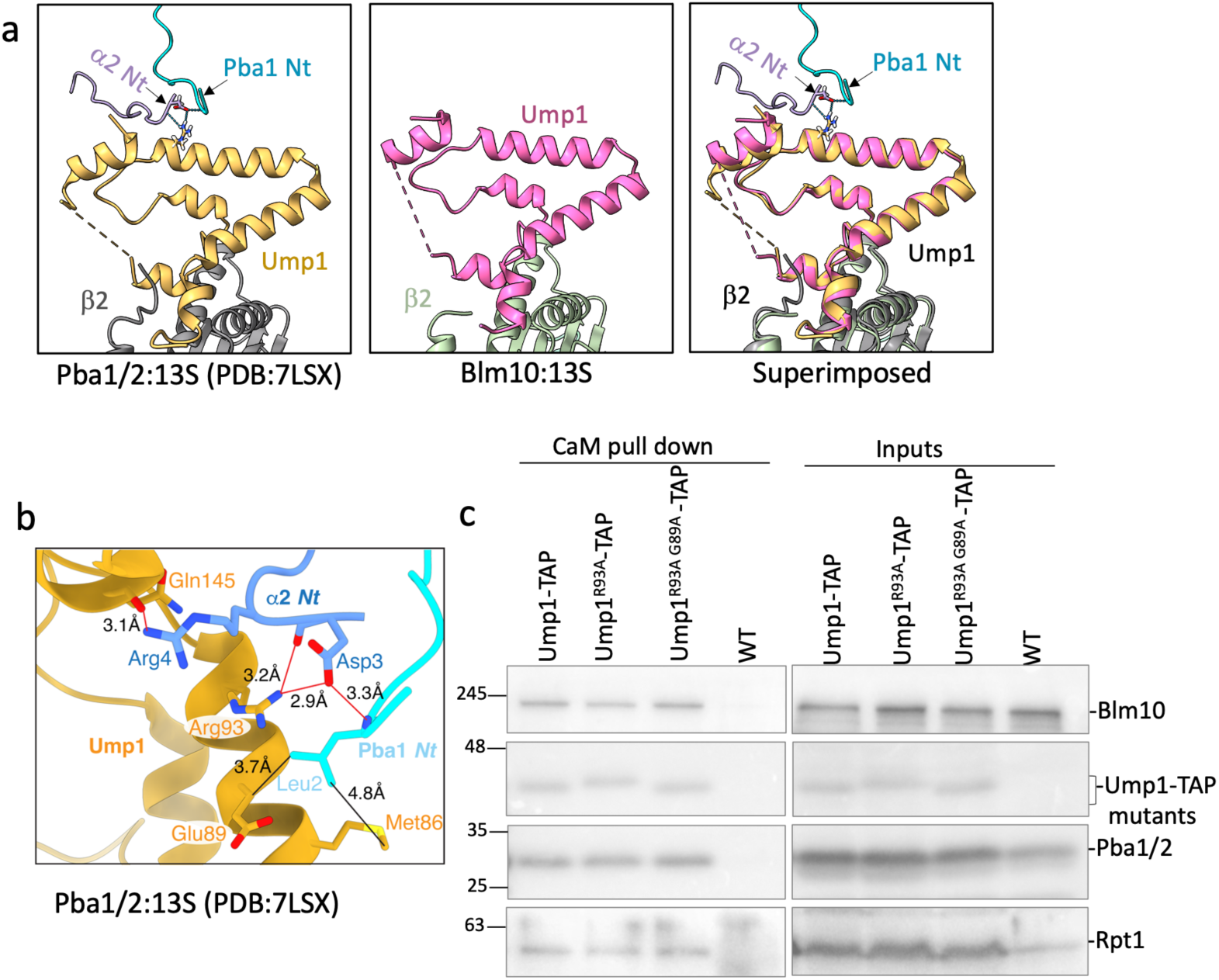
Disruption of putative Pba1/Ump1 interaction does not disrupt Pba1 binding to Pba1/2:13S. a, The position of Ump1 in Pba1/2:13S and Blm10:13S is unchanged. Panel a shows Ump1 (yellow) interacting with N-termini of Pba1 (cyan) and α2 (purple). Blue dotted lines show hydrogen bonding. The middle panel shows Ump1 (pink) alignment in Blm10:13S and the right-most panel shows the superimposed Pba1/2:13S (PDB:7LSX) and Blm10:13S to highlight the unchanged Ump1 position. **b,** The residues involved in the interaction between N-terminus of Pba1 and α2 subunit, and Ump1 in Pba1/2:13S (PDB: 7LSX) are shown. Red lines represent hydrogen bond interactions and black lines represent hydrophobic interactions with distances displayed in Å. **c,** Western blot analysis of Ump1-TAP (CaM) based pull down from mutant strains Ump1^R93A^ and Ump1^R93A^ ^E89A^ and the corresponding inputs.

**Supplementary Table 1.** Strain list.

| Strain | Genes manipulated | Repair DNA | Fig. | Ref. |
| --- | --- | --- | --- | --- |
| sUB61 | *MATα lys2-801 leu2-3, 2-112 ura3-52 his3D200 trp1-1* | n/a | 7,8b-c, 6S | (a) |
| sJR785 | *MAT A ump1::UMP1-CBP-TEV-ZZ-(His3MX6)* | n/a | 7,8b-c, 6S | (b) |
| sJR792 | *MAT A pba1::PBA1 (NAT) ump1::UMP1- CBP-TEV-ZZ-(His3MX6)* | n/a | 8c | (c) |
| sJR793 | *MAT A pba1::PBA1 (HYG) blm10::BLM10 (NAT) ump1::UMP1- CBPTEV-ZZ-(His3MX6)* | n/a | 8c | (c) |
| sJR794 | *MAT A blm10::BLM10 (NAT) ump1::UMP1- CBPTEV-ZZ-(His3MX6)* | n/a | 8c | (c) |
| sJR795 | *MAT A pba1::PBA1Δ3C (KAN) ump1::UMP1- CBPTEV-ZZ-(His3MX6)* | n/a | 7 | (c) |
| sJR809 | *MAT A pba1::PBA1 Δ3C (KAN) pba2:: PBA2 Δ3C (NAT) ump1::UMP1- CBPTEV-ZZ-(His3MX6)* | n/a | 7 | (c) |
| sJR2578 | *MAT A pba1::PBA1<sup>L2S</sup>Δ3-17 ump1::UMP1-CBP-TEV-ZZ-(His3MX6)* | pRL1478 | 7 | This study |
| sJR2583 | *MAT A pba1::PBA1<sup>L2S</sup>Δ3-31 ump1::UMP1-CBP-TEV-ZZ-(His3MX6)* | pRL1503 | 7 | This study |
| sJR2584 | *MAT A pba1::PBA1<sup>L2S</sup>Δ3-31 pba1::PBA1Δ3C (KAN) ump1::UMP1-CBP-TEV-ZZ-(His3MX6)* | pRL1503 | 7 | This study |
| sJR2586 | *MAT A pba1::PBA1<sup>L2S</sup>Δ3-31 pba1::PBA1Δ3C (KAN) pba2:: PBA2 Δ3C (NAT) ump1::UMP1-CBP-TEV-ZZ-(His3MX6)* | pRL1503 | 7 | This study |
| sJR2588 | *MAT A pba1::PBA1<sup>L2S</sup>Δ3-17 blm10::GPDpGFPBLM10 (CloNAT) ump1::UMP1-CBP-TEV-ZZ-(His3MX6)* | pRL1478 | 7 | This study |
| sJR2589 | *MAT A pba1::PBA1<sup>L2S</sup>Δ3-31 blm10::GPDpGFPBLM10 (CloNAT) ump1::UMP1-CBP-TEV-ZZ-(His3MX6)* | pRL1503 | 7 | This study |
| sJR1012 | *MAT A blm10::GPDpGFPBLM10 (NAT) ump1::UMP1-CBP-TEV-ZZ-(His3MX6)* | n/a | 7 | (d) |
| sJR2614 | *MAT A pba4::PBA4-CBP-TEV-ZZ-(His3MX6)* | n/a | 8b | (b) |
| sJR2543 | *MAT A ump1::UMP1<sup>R93A</sup> ump1::UMP1- CBPTEV-ZZ-(His3MX6)* | pRL1470 | 6S | This study |
| sJR2545 | *MAT A ump1::UMP1<sup>R93A, G89A</sup> ump1::UMP1- CBPTEV-ZZ-(His3MX6)* | pRL1471 | 6S | This study |
| **genotype is: MAT A his3Δ0 leuΔ0 met15Δ0 ura3Δ0**, except for sUB61 which is *MATα lys2-801 leu2-3, 2-112 ura3-52 his3D200 trp1-1*. (a) Finley, D., Ozkaynak, E. & Varshavsky, A. **1987** Cell 48, 1035-1046. (b) Ghaemmaghani, S. et al. **2003** Nature 425, 737-741. (c) Wani, P. et al. **2015** Nature Communications 7384, 1-11. (d) Burris, A. et al 2021 Journal of Biological Chemistry, 296, 1-18. |  |  |  |  |

**Supplementary Table 2.**
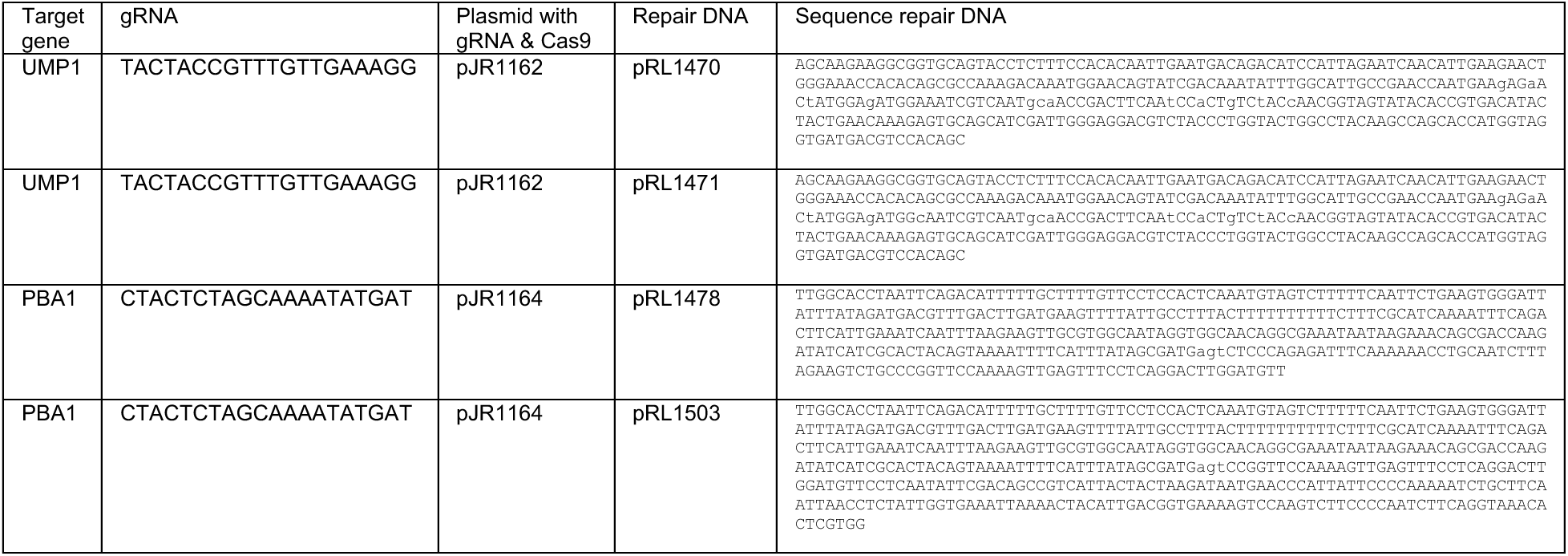
Crispr/Cas9 gRNA and repair DNA.

## Notes

### Competing Interest Statement

The authors have declared no competing interest.

